# Single-nucleus RNA sequencing of human periventricular white matter in vascular dementia

**DOI:** 10.1101/2024.12.06.627202

**Authors:** Sebastián Díaz-Pérez, Jonathan H. DeLong, Cyprien A. Rivier, Chia-Yi Lee, Michael H. Askenase, Biqing Zhu, Le Zhang, Kristen J. Brennand, Andrew J. Martins, Lauren H. Sansing

## Abstract

Vascular dementia (VaD) refers to a variety of dementias driven by cerebrovascular disease and is the second leading cause of dementia globally. VaD may be caused by ischemic strokes, intracerebral hemorrhage, and/or cerebral small vessel disease, commonly identified as white matter hyperintensities on MRI. The mechanisms underlying these white matter lesions in the periventricular brain are poorly understood. In this study we perform an extensive transcriptomic analysis on human postmortem periventricular white matter lesions in patients with VaD with the goal of identifying molecular pathways in the disease. We find increased cellular stress responses in astrocytes, oligodendrocytes, and oligodendrocyte precursor cells as well as transcriptional and translational repression in microglia in our dataset. We show that several genes identified by GWAS as being associated with white matter disease are differentially expressed in cells in VaD. Finally, we compare our dataset to an independent snRNAseq dataset of PVWM in VaD and a scRNAseq dataset on human iPSC-derived microglia exposed to oxygen glucose deprivation (OGD). We identify the increase of the heat shock protein response as a conserved feature of VaD across celltypes and show that this increase is not linked to OGD exposure. Overall, our study is the first to show that increased heat shock protein responses are a common feature of lesioned PVWM in VaD and may represent a potential therapeutic target.

## Introduction

Vascular dementia (VaD) is an umbrella term that describes dementias whose etiology is attributed to cerebrovascular pathology (1). VaD diagnosis is achieved by assessing a combination of clinical criteria, neuroimaging findings and cognitive testing. A definitive diagnosis of pure VaD is achieved postmortem by confirming the absence of co-existing non-vascular pathologies that could explain cognitive decline. In some cases, other dementias, such as Alzheimer’s disease (AD), can occur alongside cerebrovascular dysfunction (2) and, in these cases, cerebrovascular disease (CVD) has been shown to exacerbate cognitive dysfunction independently of amyloid beta burden (3). Given the heterogeneity of VaD, a classification system with six different VaD subtypes, including mixed dementia, has been proposed (4–6). Of these, cerebral small vessel disease (CSVD) is the strongest contributor progressive vascular dementia, and is identified on neuroimaging by white matter hyperintensities in the periventricular and deep subcortical white matter, lacunar infarctions, enlarged perivascular spaces, cerebral microbleeds, and brain atrophy (7).

Given the lack of local energy reserves, the brain is highly dependent on the local cerebral vasculature to meet shifting local energy demands (8). Proper modulation of blood flow to meet local energy demands is achieved through a complex process known as hemodynamic coupling (9, 10), which involves the concerted actions of multiple cell types including neurons, astrocytes, vascular endothelial and mural cells, smooth muscle cells, pericytes and microglia. Hemodynamic coupling is impaired in CSVD leading to ischemic lesions in the periventricular and deep white matter of the brain, areas which are particularly vulnerable to hypoperfusion (11–15). These lesions are characterized pathologically by demyelination, mild gliosis, enlarged perivascular spaces, and tissue edema (16). However, pathological data distinguishing periventricular and deep white matter lesions is limited. Over time, progression of this CSVD-driven pathology can result in the development of VaD or exacerbation of mixed dementia (3). Our mechanistic understanding of VaD pathology remains incomplete and there is currently no treatment that can cure, reverse or halt the progression of VaD (17).

Several studies have linked neuroinflammation to the progression of other neurodegenerative diseases like AD (18–20). In the case of VaD, there is evidence from studies of postmortem tissue that support neuroinflammation playing a role in disease. Neuropathological studies of VaD brains have shown demyelination, reactive gliosis (21, 22) and hypoxia-inducible factor 1 alpha (HIF1A) activation (23) at the site of white matter lesions. Postmortem transcriptomic analysis showed that deep subcortical lesions and the surrounding white matter had upregulation of pro-inflammatory genes *C1QTNF3*, *CD14*, *HLA-DQB1* and *ISG15* (24). Finally, single-nucleus RNA sequencing (snRNAseq) at the lesion sites in periventricular white matter (PVWM) has shown increased pro-inflammatory gene expression in microglia (25).

Additional evidence comes from rodent models of chronic cerebral hypoperfusion (CCH) (26). CCH is usually achieved by insertion of microcoils or ameroid constrictor rings around the common carotid arteries which leads to a reduction of cerebral blood flow. These models show some pathological features similar to VaD including long-term reactive gliosis, loss of myelin density and impaired cognitive performance (27–31). While promising, it is important to note that these models also suffer from key pitfalls. Acute occlusion or stenosis of the carotid arteries during surgery leads to acute reduction of cerebral blood flow (CBF) and subsequent compensation of CBF over time (32), a phenomenon that may be more reflective of acute, moderate ischemia and not the gradual reduction of CBF in human VaD in the microvasculature. CCH also does not specifically target the white matter, and several CCH models show injury of both white and grey matter. Finally, the primarily focus of most CCH studies has been on the first 30 days post-surgery with less data on later timepoints. Current studies with CCH models have shown CCH induces a chronic increase of IL-1β, TNF and IL-6 (33–37) while inhibition of pro-inflammatory mediators like HMGB1 (37), AIM2 (38) and IL-1β (39), has led to decreased white matter injury and improved cognition by day 28 post surgery. Taken together, evidence suggest that neuroinflammation may play a role in VaD pathology.

While significant progress has been made, the heterogeneity of disease (1, 4, 5) and the limitations of current animal models, have hindered our ability to gain mechanistic insight into VaD. To address this gap, we focus on studying CSVD-driven disease using snRNAseq on human post-mortem PVWM from healthy controls and VaD patients. We perform a comprehensive transcriptomic analysis involving differentially expressed genes (DEGs), gene set enrichment analyses (GSEA), predicted receptor-ligand interaction analysis and network analysis. We then use Rank-Rank Hypergeometric Analysis (RRHO) and module scoring to assess the overlap between our dataset and a previously published snRNAseq dataset on VaD (25) in order to identify shared signatures across independent patient cohorts. We also test the extent to which exposure of human iPSC-derived microglia (iMGL) to oxygen-glucose deprivation (OGD) recapitulates the transcriptional signature seen in VaD. Finally, we use Multi-marker Analysis of GenoMic Annotation (MAGMA) to test whether any of the genes identified by genome-wise association studies (GWAS) to be associated with CSVD, AD and ICH were overexpressed in any of the cell-type transcriptomic signatures identified in our dataset. Overall, we describe an abundance of cell stress responses in the transcriptomic signatures in our dataset, we show that exposure of iMGL to OGD is capable of recapitulating key aspects of the transcriptomic signature observed in VaD microglia, we show the upregulation of the heat shock protein (HSP) response as a conserved signature in VaD and show significant association of CSVD genetic risk variants to our transcriptomic signatures in VaD.

## Results

### Characterizing the cellular transcriptome of periventricular white matter in VaD

In order to determine the cellular transcriptomic profiles in PVWM in VaD versus controls, we performed single-nucleus RNA sequencing on periventricular brain tissue adjacent to the left frontal horn (Figure 1A). Healthy and VaD groups were matched for age and sex (age range 61-100 years, 2 males and 2 females per group) (Supplemental Table 1). To exclude cases where AD was the primary contributor of dementia, only samples with Braak & Braak staging II or lower were selected for processing. Four brains in each group passed our quality metrics and were included in the analysis (Supplementary Table 2) (5). Astrocytes, microglia, oligodendrocytes, oligodendrocytes precursor cells (OPCs), neurons and endothelial cells were identified based on expression of cell type-specific markers (Figure 1B, C and D, Supplementary Table 3). As expected with white matter tissue, and as previously observed (25), the majority of cells were oligodendrocytes while neurons and endothelial cells were relatively scarce (Figure 1D). No significant differences in the proportions of each cell type were observed between control and VaD samples.

**Figure 1.**
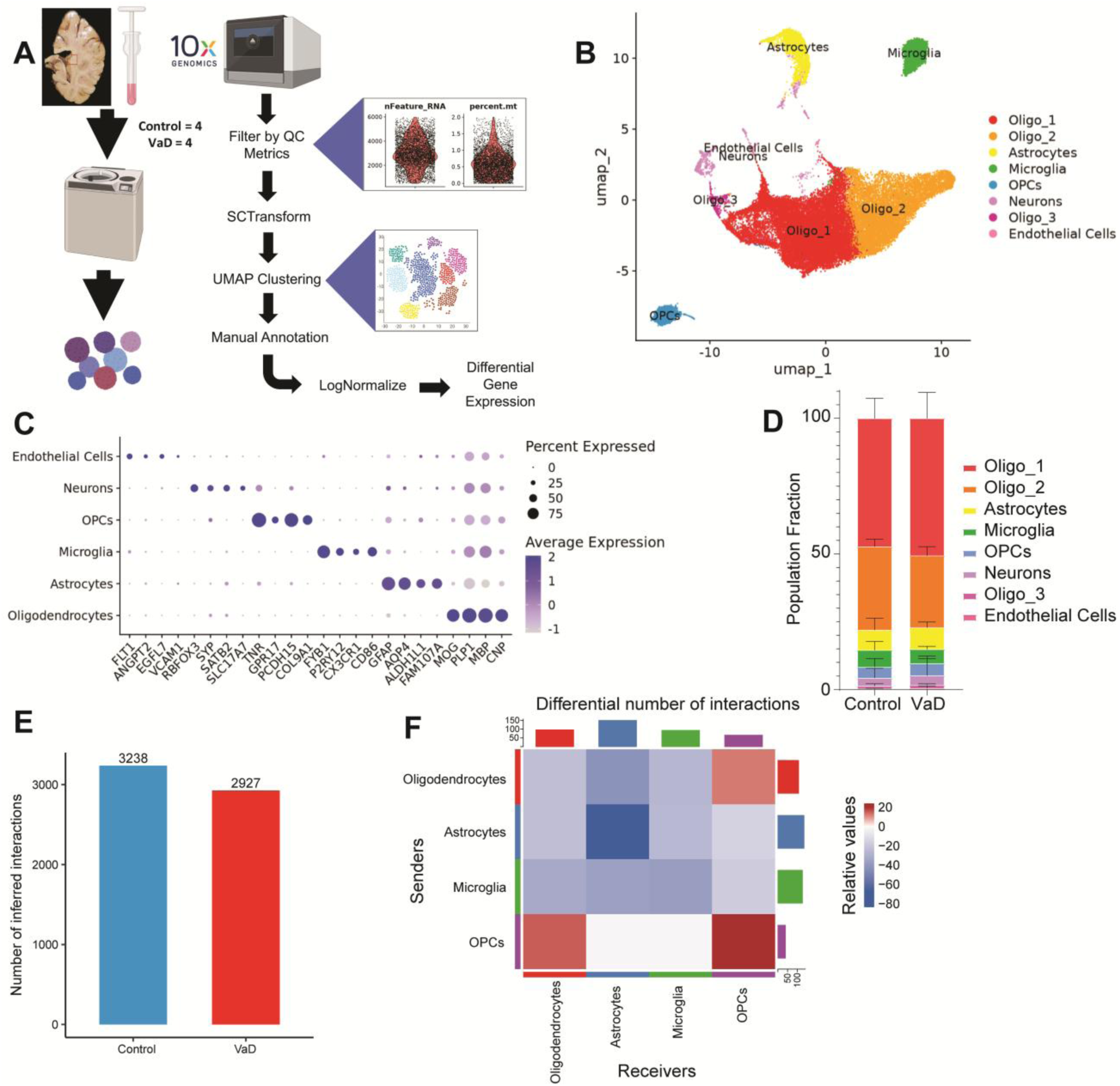
snRNAseq of human post-mortem periventricular white matter. (**A**) Diagram summarizing the steps used to create our snRNAseq dataset. (**B-C**) UMAP clustering of snRNAseq dataset annotated based on expression of cell type-specific markers. (**D**) Proportion of cell types found in Control vs VaD samples. (**E-F**) Overview of differential receptor ligand (R-L) signaling across cell types in VaD. N=4 brain samples/group.

Because of their low numbers, we excluded neurons and endothelial cells from further analysis. Using CellChatDB, we identified the total number of predicted receptor-ligand interactions found across cell types in control and VaD samples (Figure 1, E and F). We observed an overall decrease in the total number of predicted receptor-ligand interactions in VaD samples. Looking at these interactions by cell type we observed a small increase only in interactions involving OPCs and oligodendrocytes, whereas cell-cell interactions were decreased between astrocytes, microglia and oligodendrocytes in VaD samples.

### Cellular stress underlies the astrocyte transcriptome in VaD samples

To determine differentially enriched biological pathways in astrocytes between subjects with VaD and controls we performed GSEA. We observed a marked upregulation of pathways related to cadherin production, HSP responses, cholesterol signaling and NRAGE-mediated apoptosis (Figure 2A). To understand the broader contours of the astrocyte transcriptome, we used a less stringent filter (FDR ≤ 0.25), as previously described (40), to select pathways used to perform network analysis using the Cytoscape EnrichmentMap plug-in (Figure 2B). This approach groups GSEA-derived pathways by gene overlap, which can then be further annotated (see Methods), allowing us to understand the broader biological functions that may be dysregulated in VaD. Using network analysis, we see the main biological functions upregulated in VaD astrocytes include cell adhesion, HSP responses and p75NTR-mediated apoptosis. We also found increased cellular senescence and increased histone activity (driven by expression of histone deacetylases (HDACs) and histone demethylases). While we did not observe any downregulated pathways via GSEA, there were several receptor-ligand interactions involving astrocytes found to be downregulated. These included interactions related to TAM receptor-mediated efferocytosis, PECAM1-mediated cell adhesion, production of macrophage migration inhibitory factor (MIF), production of apolipoprotein E (APOE) and production of angiopoietins (ANGPT) (Figure 2, C and D).

**Figure 2.**
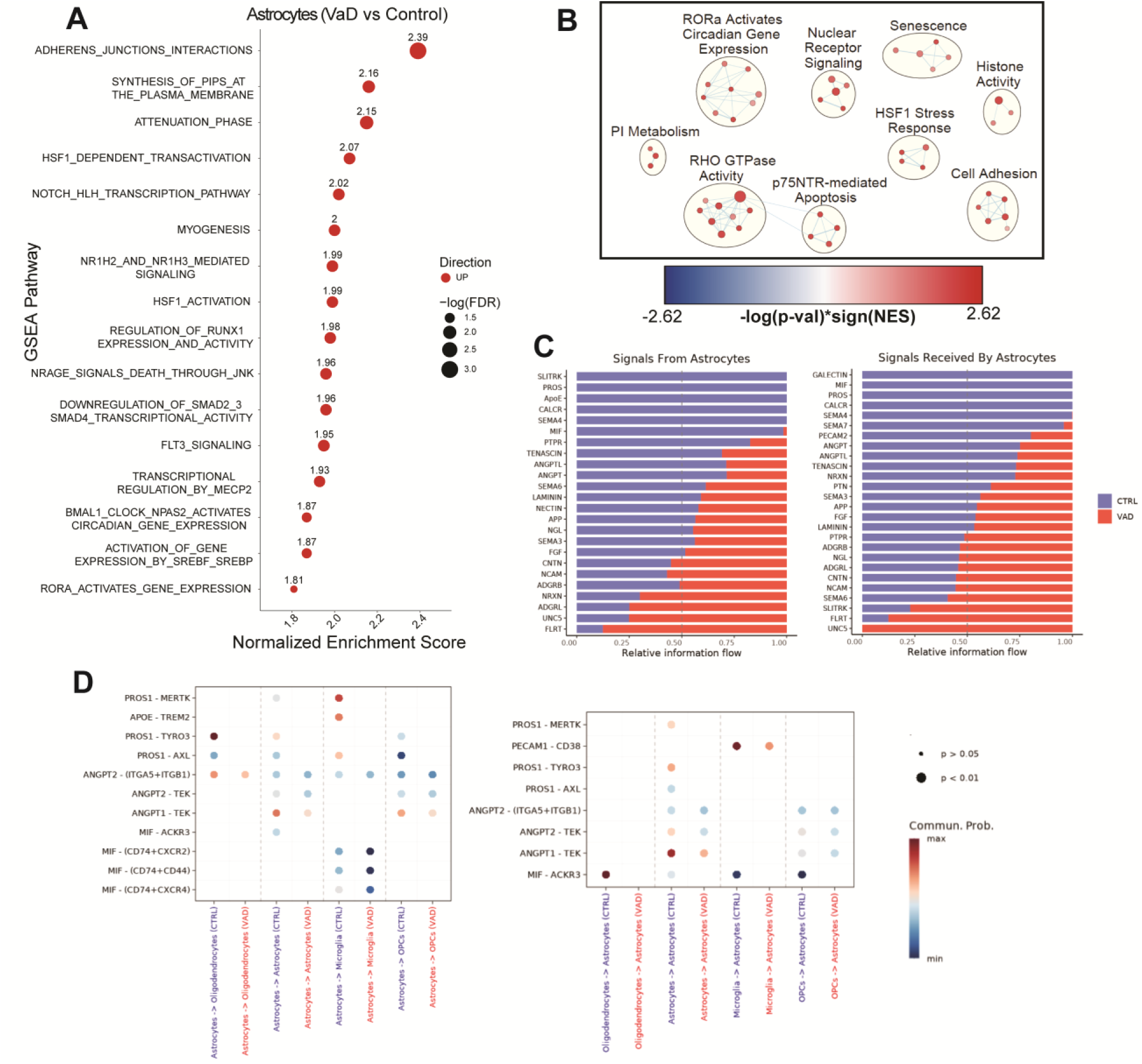
Increased cell adhesion and heat shock protein responses characterize the astrocyte transcriptome in VaD. (**A**) GSEA pathways differentially regulated in VaD astrocytes (FDR ≤ 0.05). (**B**) Cytoscape EnrichmentMap showing individual GSEA hits (FDR ≤ 0.25) clustered via AutoAnnotate. (**C-D**) Pathways and specific R-L interactions predicted by CellChatDB to be differentially regulated in VaD astrocytes.

### Impaired cellular functions underlie the transcriptome of VaD microglia

Following the same analytical approach employed for astrocytes, we analyzed the transcriptomes of microglia, oligodendrocytes and OPCs. The top upregulated pathway in VaD microglia is transcriptional repression by HDACs (“Reactome HDACS Deacetylate Histones” R-HSA-3214815) while most downregulated pathways relate to cellular translation and transcription (Figure 3A). This downregulation of transcription and translation coupled to increase HDAC activity was also observed when performing Cytoscape network analysis (Figure 3B). Despite the global downregulation of transcription and translation, the network analysis also showed increases in cell adhesion and cholesterol biosynthesis. Receptor-ligand interaction analysis showed a decrease in interactions related to key microglial functions including decreased TAM receptor-mediated efferocytosis, APOE binding, APP binding, galectin-9 production (LGALS9) and MIF signaling (Figure 3, C and D).

**Figure 3.**
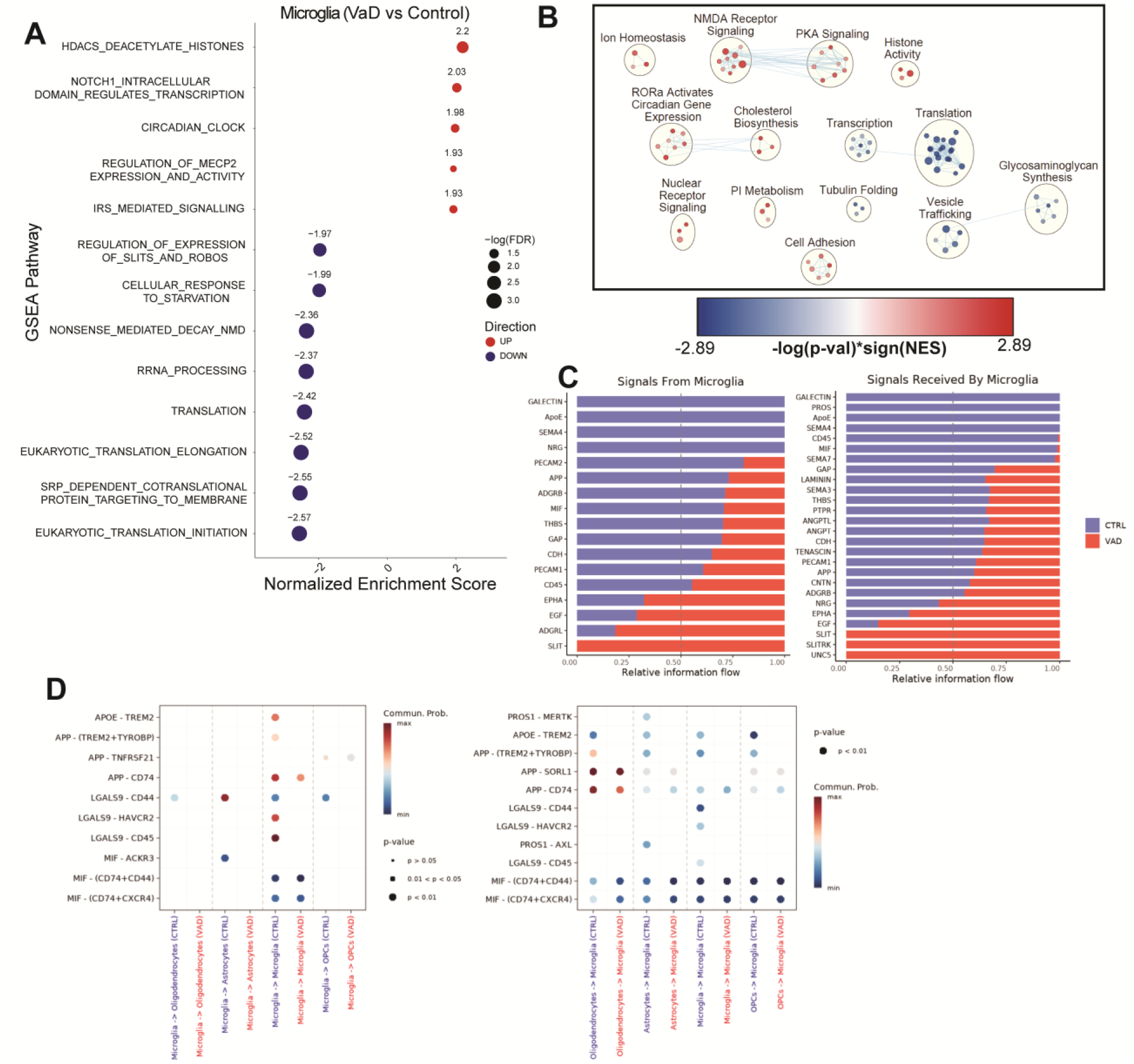
Transcriptional and translational repression characterize the microglial transcriptome in VaD. (**A**) GSEA pathways differentially regulated in VaD microglia (FDR ≤ 0.05). (**B**) Cytoscape EnrichmentMap showing individual GSEA hits (FDR ≤ 0.25) clustered via AutoAnnotate. (**C-D**) Pathways and specific R-L interactions predicted by CellChatDB to be differentially regulated in VaD microglia.

### Compromised myelination, cellular stress and increased cholesterol biosynthesis underlie the oligodendrocyte transcriptome in VaD

GSEA analysis of VaD oligodendrocytes showed altered cellular metabolism, including an upregulation of cholesterol biosynthesis and a downregulation of oxidative phosphorylation (Figure 4A). Cytoscape network analysis largely highlights the decrease in oxidative phosphorylation and increased cholesterol biosynthesis. It also reveals a cellular stress signature characterized by a downregulation of cellular transcription, protein folding and vesicle trafficking (Figure 4B). Finally, several receptor-ligand interactions implicated in myelination, such as those mediated by myelin-associated glycoprotein (MAG), claudin 11 (CLDN11) and neuregulins (NRG), were found to be decreased in VaD oligodendrocytes (Figure 4, C and D).

**Figure 4.**
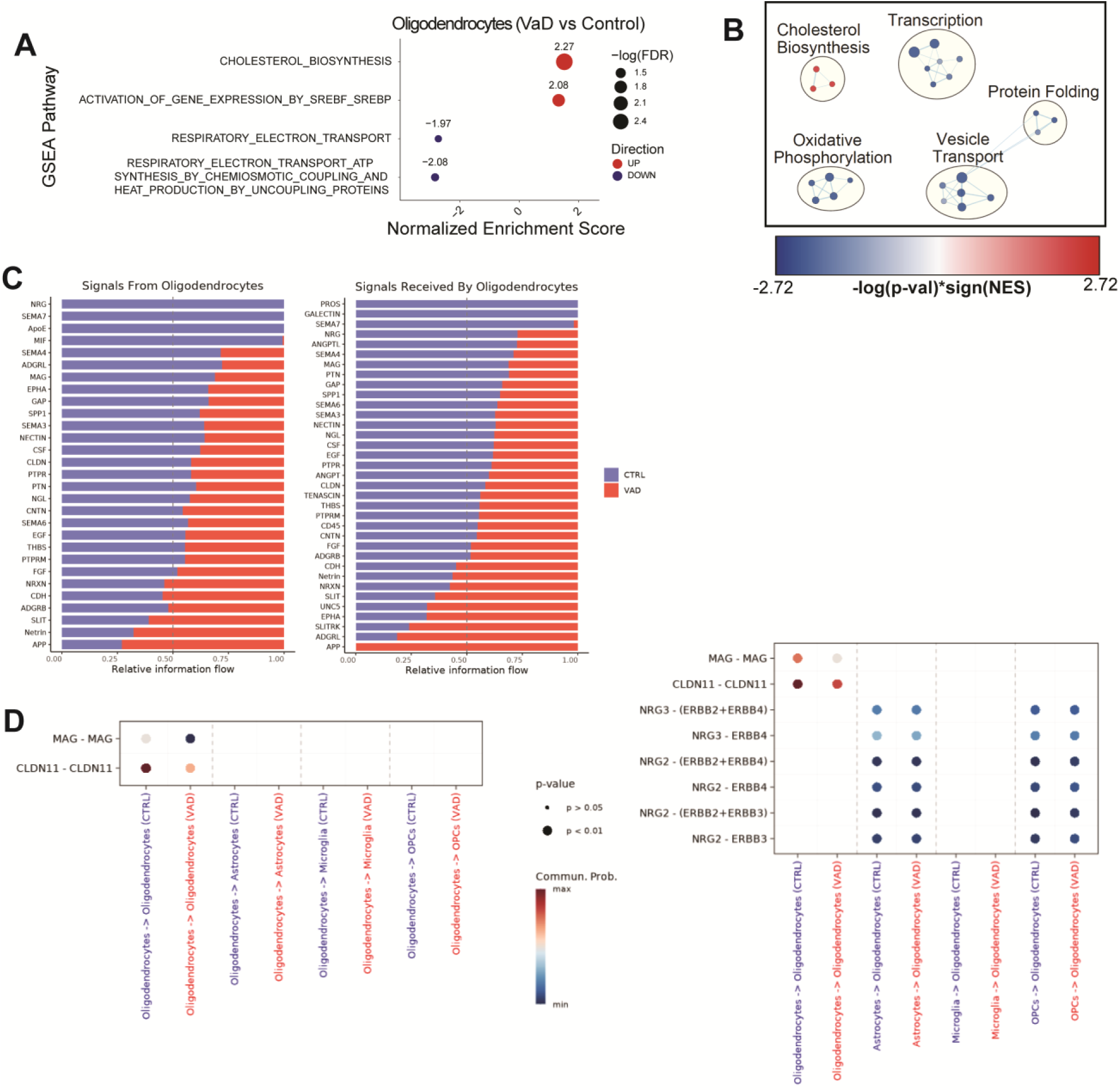
Increased cholesterol biosynthesis and impaired oxidative phosphorylation characterize the oligodendrocyte transcriptome in VaD. (**A**) GSEA pathways differentially regulated in VaD oligodendrocytes (FDR ≤ 0.05). (**B**) Cytoscape EnrichmentMap showing individual GSEA hits (FDR ≤ 0.25) clustered via AutoAnnotate. (**C-D**) Pathways and specific R-L interactions predicted by CellChatDB to be differentially regulated in VaD oligodendrocytes.

### Cellular stress responses and attempts at remyelination underlie the OPC transcriptome in VaD

GSEA analysis of VaD OPCs mainly showed an upregulation of HSP responses (Figure 5A), similar to VaD astrocytes. Cytoscape Network Analysis shows additional biological functions upregulated in VaD OPCs including NMDA receptor signaling as well as other signaling cascades with broad biological functions like phosphatidylinositol (PI), protein kinase A (PKA), RHO GTPases and nuclear receptors (Figure 5B).

**Figure 5.**
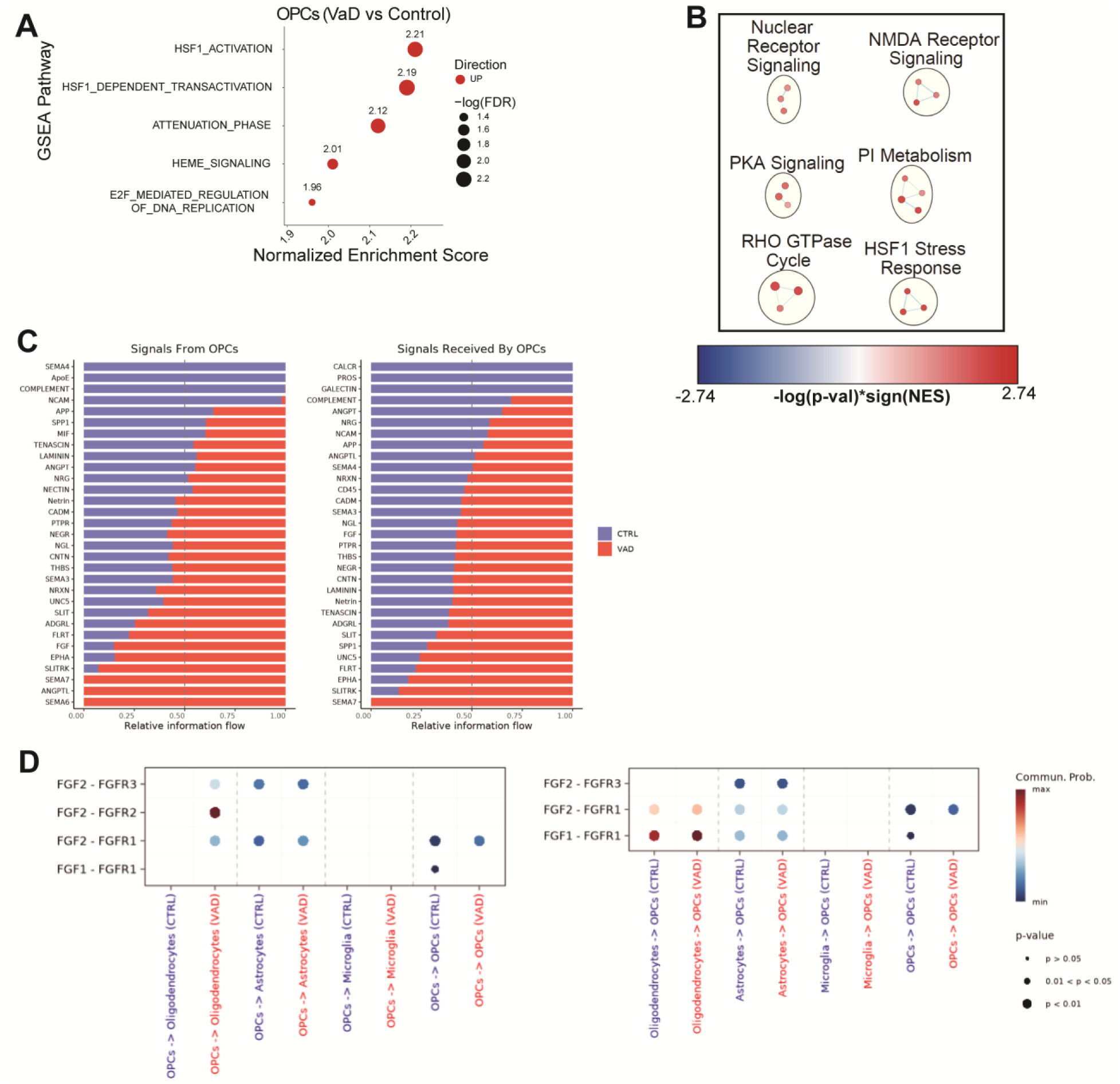
Increased heat shock protein responses characterize the oligodendrocyte precursor cell (OPCs) transcriptome in VaD. (**A**) GSEA pathways differentially regulated in VaD OPCs (FDR ≤ 0.05). (**B**) Cytoscape EnrichmentMap showing individual GSEA hits (FDR ≤ 0.25) clustered via AutoAnnotate. (**C-D**) Pathways and specific R-L interactions predicted by CellChatDB to be differentially regulated in VaD OPCs.

While OPCs are well-known for their role in promoting myelination by differentiating into myelinating oligodendrocytes, OPCs have also been shown to participate in several other non-myelinating functions such as angiogenesis, mediating neuroinflammation and regulating neuronal migration (41). Several receptor-ligand interactions related to these functions were identified by CellChat as differentially regulated in VaD OPCs. Some of the top receptor-ligand hits enriched in VaD OPCs are related to axon guidance and repulsion in the central nervous system (CNS) including EPHA, FLRT, SLIT, SLITRK, semaphorins, and UNC5 (Figure 5C). While interesting, these interactions must be interpreted with caution given the lack of a robust neuronal population in our dataset. FGF signaling is known to promote proliferation and survival of various cells in the CNS (42). Diving into the individual receptor-ligand interactions from FGF signaling, we observed an overall increased in FGF1 and FGF2-mediated signaling in subjects with VaD (Figure 5D). Both FGF1 and FGF2-mediated signaling has been shown to be pivotal for oligodendrocyte maturation and myelination (42). It’s possible that an increase in FGF signaling may be an attempt to promote maturation and remyelination at the lesion site.

### Cellular stress responses do not correlate with increased post-mortem interval

Degradation of mRNA transcripts can occur quickly, raising the possibility that long post-mortem intervals (PMI) may affect the quality of the transcriptome in *postmortem* samples (43). It has been previously shown that when samples are stored under appropriate conditions, RNA can persist largely intact up to 48 hrs postmortem (44), which is above the highest PMI amongst our samples (30 hrs). However, cognizant of the sizeable difference in the average PMI between controls (7.65 hrs) and VaD (18.41 hrs) (Supplementary Table 1), we sought to test whether increasing PMI may be driving some of the transcriptomic differences observed in our dataset. Because increased HSP responses were a conserved feature in our dataset, we used a Pearson Correlation Coefficient test to see whether increasing PMI correlated with increased expression of genes involved in the HSP response (Supplementary Table 4). Overall, regardless of whether the gene expression was averaged for the entire dataset or by cell type, we did not observe significant associations between increasing PMI and HSP response gene expression.

### Cellular stress responses represent a limited overlap between two VaD transcriptomic datasets

Transcriptomic analyses have the potential to notably advance our understanding of VaD pathology. In 2022, Mitroi et al published a single-nucleus RNA sequencing study on VaD also focusing on PVWM (25). While the protocols used in their study and ours are largely similar, one notable difference was the nuclei sorting strategy. The published data set FACS-sorted their isolated nuclei prior to sequencing for either DAPI expression or DAPI and ERG expression which allowed them to enrich for endothelial cells and also identified the microglia. We did not FACS-sort nuclei prior to sequencing.

We reanalyzed their dataset and recapitulated similar clusters to those of their original publication and then performed GSEA and pathway grouping using Cytoscape (Supplementary Figure 1). To identify common genes that were differentially expressed in VaD across both datasets, we used rank rank hypergeometric overlap (RRHO), a threshold-free approach that uses continuous pre-ranked gene lists to identify genes with significant overlap (45). We performed RRHO to compare astrocytes, oligodendrocytes and OPCs from our unsorted dataset populations and their dataset’s DAPI-sorted populations, and since their DAPI-sorting approach did not yield microglia, we performed the RRHO comparison between our unsorted microglia and their ERG-sorted microglia. The number of overlapping genes identified by RRHO is illustrated in Venn diagrams (Supplementary Figure 2, A-D).

All cell types showed overlap in the bottom left quadrant of the heatmap, which represents genes that are upregulated in both datasets. Astrocytes in both datasets demonstrated upregulation of several stress responses including a broad array of pathways driven by HSPs (Figure 6A). Microglia from the datasets mainly showed an overlap in pathways related to chromatin remodeling (Figure 6B). Oligodendrocytes showed an overlap in several secondary messenger cascades such as DAG/IP3, protein kinase A (PKA), PLC Beta and calcium/calmodulin. They also showed overlap in upregulated cholesterol biosynthesis (Figure 6C). Finally, OPCs showed an overlap mostly in HSP responses (Figure 6D). Overall, RRHO analysis showed that upregulated HSP responses was a common overlapping signature across all cell types in both our dataset and the *Mitroi et al* dataset suggesting cellular stress may be a conserved feature in VaD. No downregulated pathways were overlapping across the data sets.

**Figure 6.**
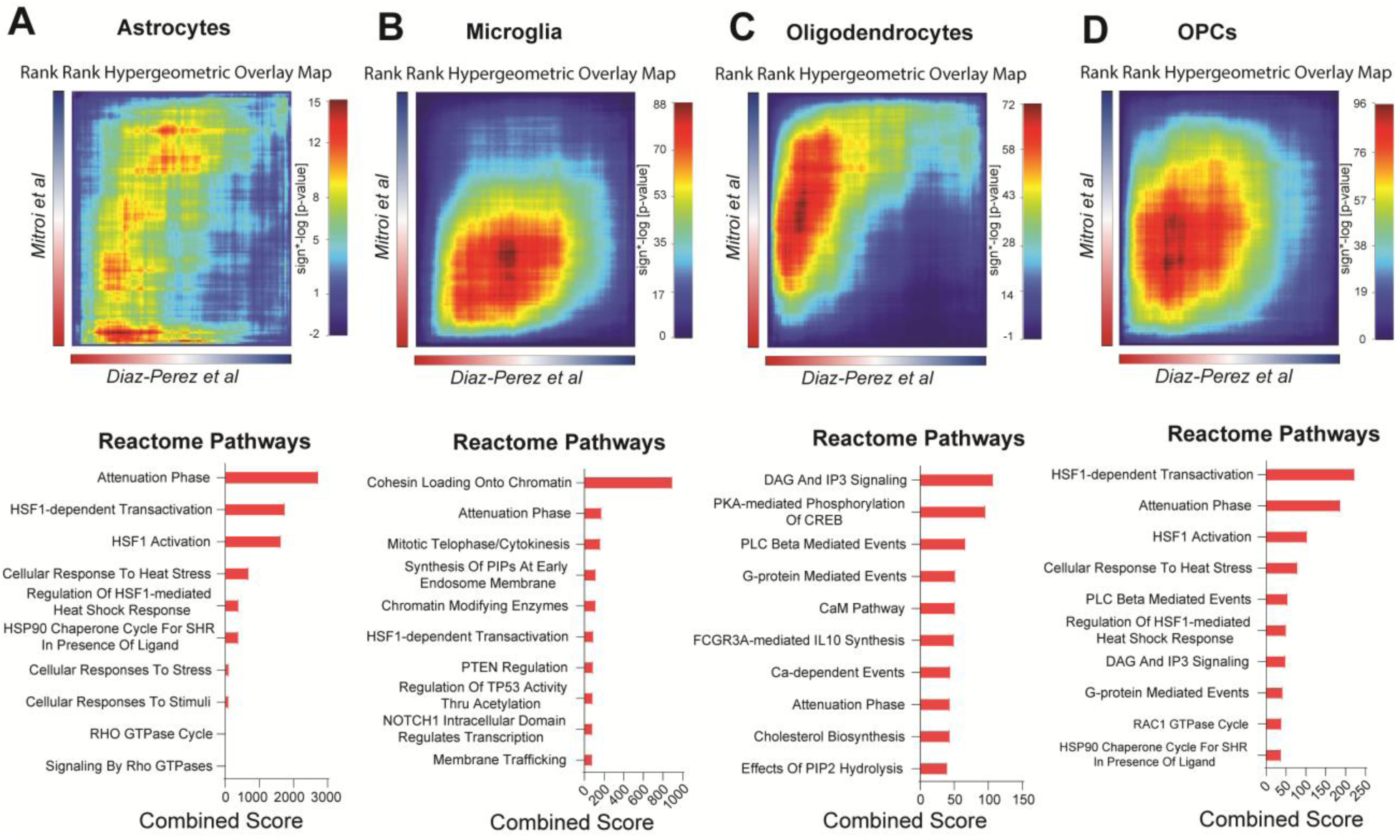
Rank-Rank Hypergeometric analysis reveals increased heat shock protein responses as a conserved feature across two independent datasets. RRHO heatmaps showing overlap of pre-ranked gene lists (VaD vs Control) from (**A**) astrocytes, (**B**) microglia, (**C**) oligodendrocytes and (**D**) OPCs from out dataset (*Diaz-Perez et al*) and the *Mitroi et al* dataset. Overlap on the top right corner of the heatmap is indicative of common downregulated genes while overlap on the bottom left is indicative of common upregulated genes. Heatmap is colored by signed −log(p-value). Top 10 Reactome gene sets from overlapping genes are shown below the respective heatmap as barplots ranked by combined score.

Both the HSP response and the unfolded protein response (UPR) are activated by proteotoxic stress. While HSPs respond to accumulation of misfolded proteins in the cytosol while the UPR responds to the accumulation of misfolded proteins in the ER (46). Given the widespread increase of HSP responses across multiple cell types in both datasets, we decided to test whether the UPR was also increased in VaD across either dataset. To do so, we tested two corresponding gene sets using Seurat’s module scoring approach (47): “HSF1 Activation” (R-HSA-3371511), and “Unfolded Protein Response UPR” (R-HSA-381119) (Figure 7, A-C). Only a subset of cells in each dataset showed a stark upregulation of the HSP response, but overall cells from the VaD group showed a global mild upregulation in the HSP response compared to control (Figure 7B). Module scoring of the UPR showed little to no difference between control and VaD, suggesting that while the HSP response is upregulated in VaD, the UPR is not (Figure 7C).

**Figure 7.**
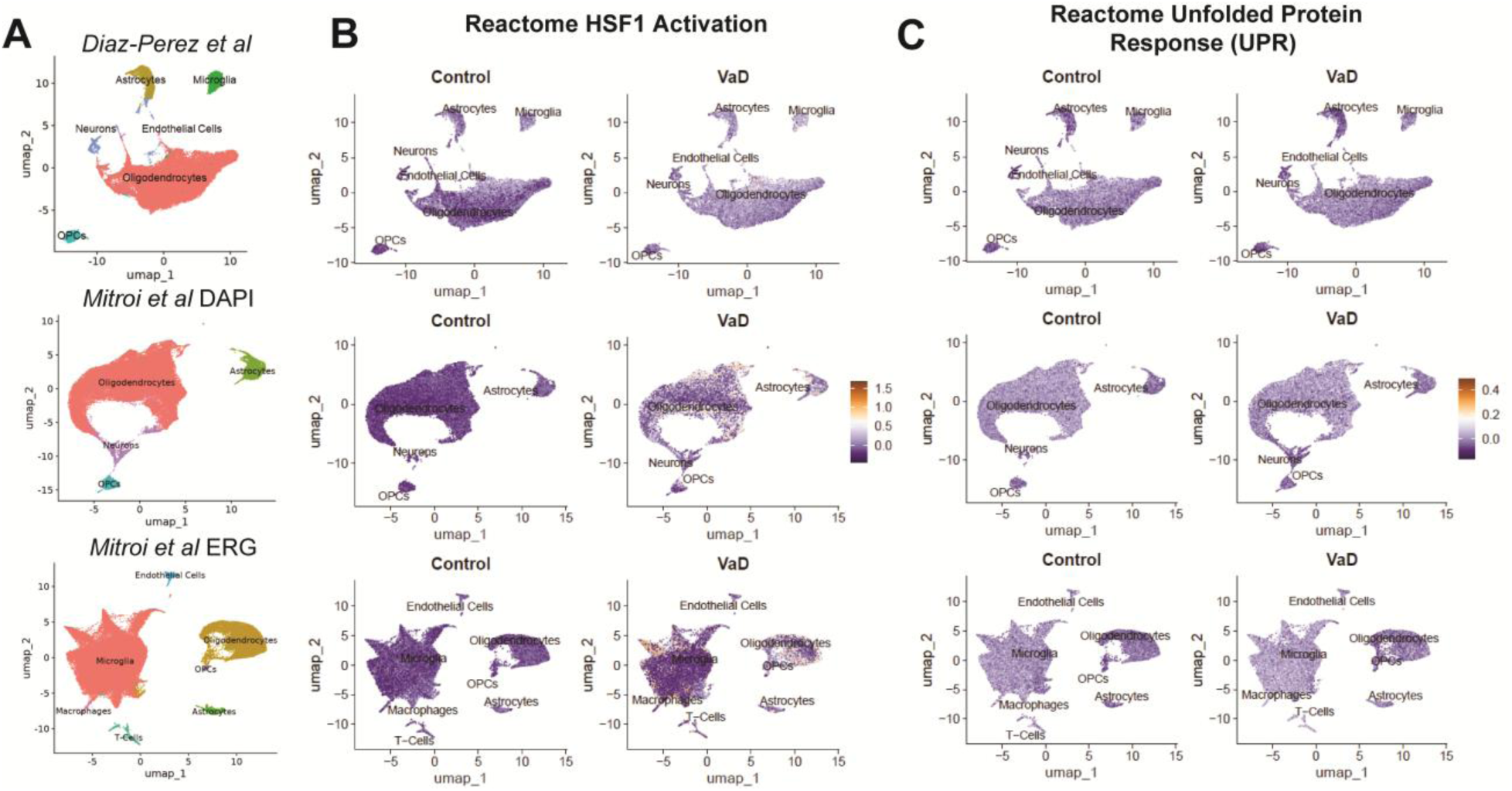
Module scoring of heat shock protein responses and unfolded protein responses across the two independent snRNAseq VaD datasets. (**A**) UMAPs representing our dataset (*Diaz-Perez et al*) and the *Mitroi et al* dataset. Module scoring of the (**B**) “Reactome HSF1 Activation” and the (**C**) “Reactome Unfolded Protein Response (UPR)” gene sets.

### Oxygen-glucose deprivation induces translational and transcriptional repression in human iPSC-derived microglia and partially recapitulates our microglial VaD transcriptomic signature

To test whether oxygen-glucose deprivation (OGD) via ischemia may induce the transcriptomic differences observed in VaD microglia, we performed an *in vitro* experiment using human microglia induced pluripotent stem cells from two donors (iMGL). We incubated differentiated iMGL cells under normoxia, hypoxia or OGD conditions for 24 hrs prior to collecting them for single-cell RNA sequencing. We observed significant heterogeneity in the cellular activation states in the iMGL cultures (Supplementary Figure 3A). All clusters showed robust expression of microglia-specific markers and the absence of stem cell, astrocyte, oligodendrocyte, OPC, endothelial cell or neuronal markers (Supplementary Figure 3B). Looking at cluster proportions, Micro_4 seemed to be preferentially depleted compared to other clusters in hypoxia and OGD (Supplementary Figure 3, C and D). Cell cycle scoring suggests that Micro_4 represents a subset of proliferating microglia that is a well-defined subcluster of cells (Supplementary Figure 3, E and F). This may suggest that these proliferating microglia are particularly susceptible to hypoxia-induced death or that hypoxia and OGD may be preferentially suppressing the transcriptomic signature that characterized this cluster.

While we observed remarkable heterogeneity in the iMGL cultures at the single cell level, IPA comparison analysis showed similar transcriptomic responses of all clusters to OGD exposure (Figure 8A), thus we proceeded to do all downstream analysis on the entire iMGL population. Using RRHO we tested whether the OGD transcriptomic signature overlapped with the microglial signature identified in the PVWM of VaD subjects (Figure 8B; Supplementary Figure 2E). Downregulated pathways involved in the translational and transcriptomic machinery of the cell were consistent across the iMGL OGD and VaD datasets (Figure 8C). In contrast, the RRHO heatmap for overlap with the *Mitroi et al* VaD dataset shows an almost overall anti-correlation with the OGD gene list and overlapping genes were limited (Supplementary Figure 4, A and B).

**Figure 8.**
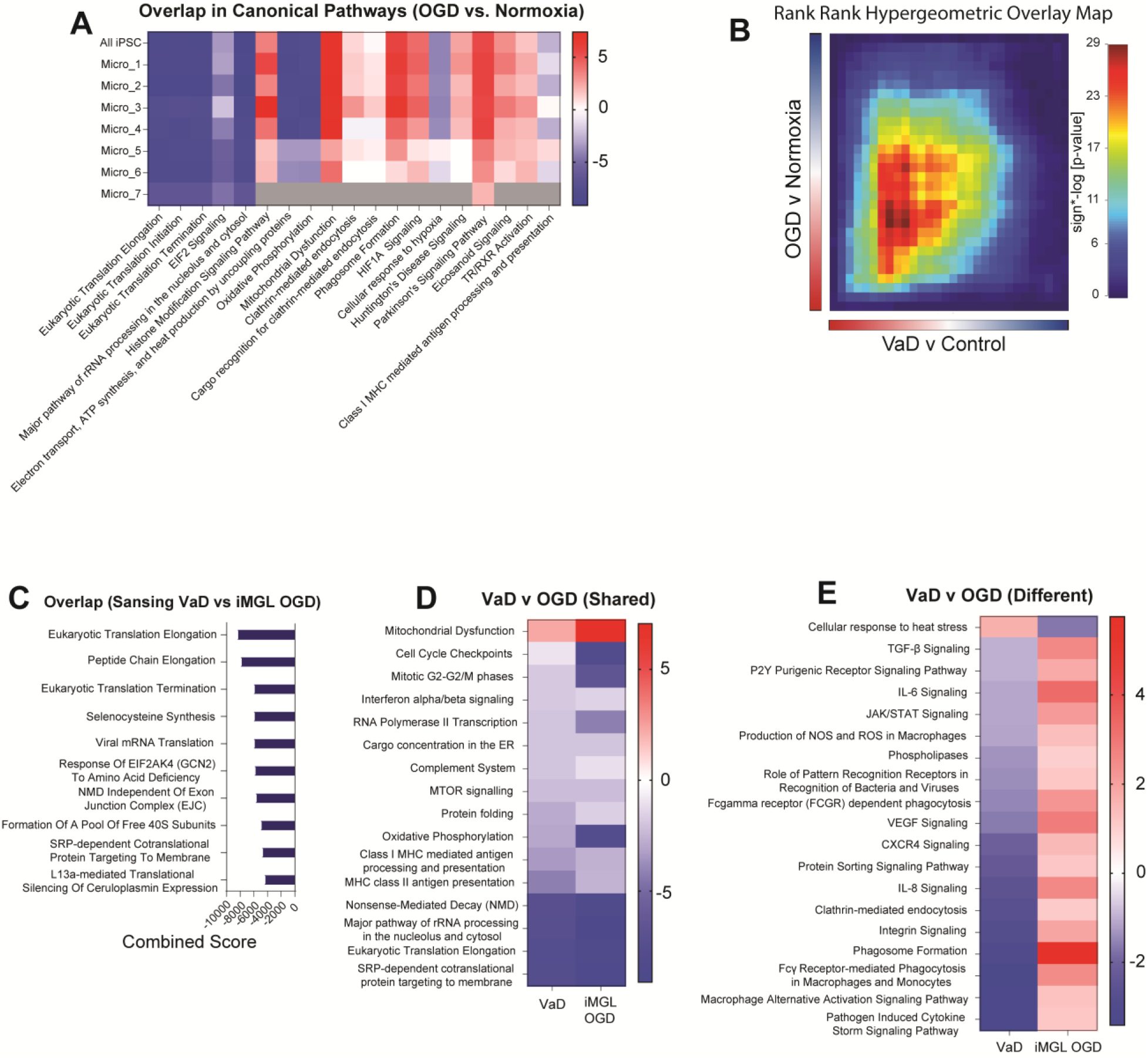
Exposure of iMGL to OGD recapitulates transcriptional and translational repression observed in VaD microglia. (**A**) IPA comparison analysis of OGD vs Normoxia in entire iMGL dataset and each individual subcluster. Gray values represents pathways that did not achieve statistical significance. (**B**) RRHO heatmap showing overlap of pre-ranked gene lists from VaD vs Control and OGD vs Normoxia. (**D**) Top 10 Reactome gene sets from overlapping genes ranked by combined score. (**E**) Selected shared and (**F**) divergent pathways obtained from comparison analysis in IPA between VaD vs Control and OGD vs Normoxia.

Because RRHO only shows us the shared transcriptomic signature between the two datasets, we employed IPA comparison analysis to determine both shared and divergent canonical pathways. The shared signature determined by IPA showed a common downregulation of the translational and transcriptomic machinery of the cell, but it also showed a downregulation of oxidative phosphorylation, complement, antigen presentation and type I IFN signaling (Figure 8D). In addition to this common signature, there were also many divergent pathways (Figure 8E). Unlike the microglial signature in VaD, exposure of iMGL to OGD induced upregulation in phagocytosis, innate immune signaling, production of reactive oxygen species (ROS) and VEGF signaling. Interestingly, as opposed to the common upregulation of HSP responses observed in VaD across datasets, iMGLs exposed to OGD show a downregulation of heat stress responses.

### Multi-marker Analysis of GenoMic Annotation

Several robust GWAS of phenotypes associated with CSVD have been published (Supplementary Table 5). Genetic risk variants identified in these studies can be mapped to specific genes making possible a cross-reference of GWAS-identified genetic risk variants and the differential expression of those genes in disease. To analyze whether any of genes associated with CSVD were overexpressed in our cell type-specific differential expression profiles from VaD, we used Multi-marker Analysis of GenoMic Annotation (MAGMA, v.1.10) (48). We used several GWAS of different CSVD phenotypes including: WMH volume (49), perivascular spaces in the white matter in European ancestry (50), small vessel stroke in European ancestry (51), and Fractional Anisotropy (FA) (52). As a negative control, we considered red hair color (53).

Out of the four CSVD-associated phenotypes that we tested, only two of them, WMH volume (Figure 9A, B and C) and fractional anisotropy (Figure 9D, E and F), showed significant association with VaD transcriptomes. Both WMH and FA are MRI-based measurements that can be used to assess white matter integrity (52) and their appearance and progression precede the onset of dementia by several years (54, 55). Meanwhile, small vessel stroke and enlarged perivascular spaces (Supplementary Figure 5, A and B), phenotypes that are not white matter-specific, did not show significant association with the transcriptomal changes in our VaD dataset. As expected, none of the VaD cell type transcriptomes were associated with red hair color (Supplementary Figure 5C).

**Figure 9.**
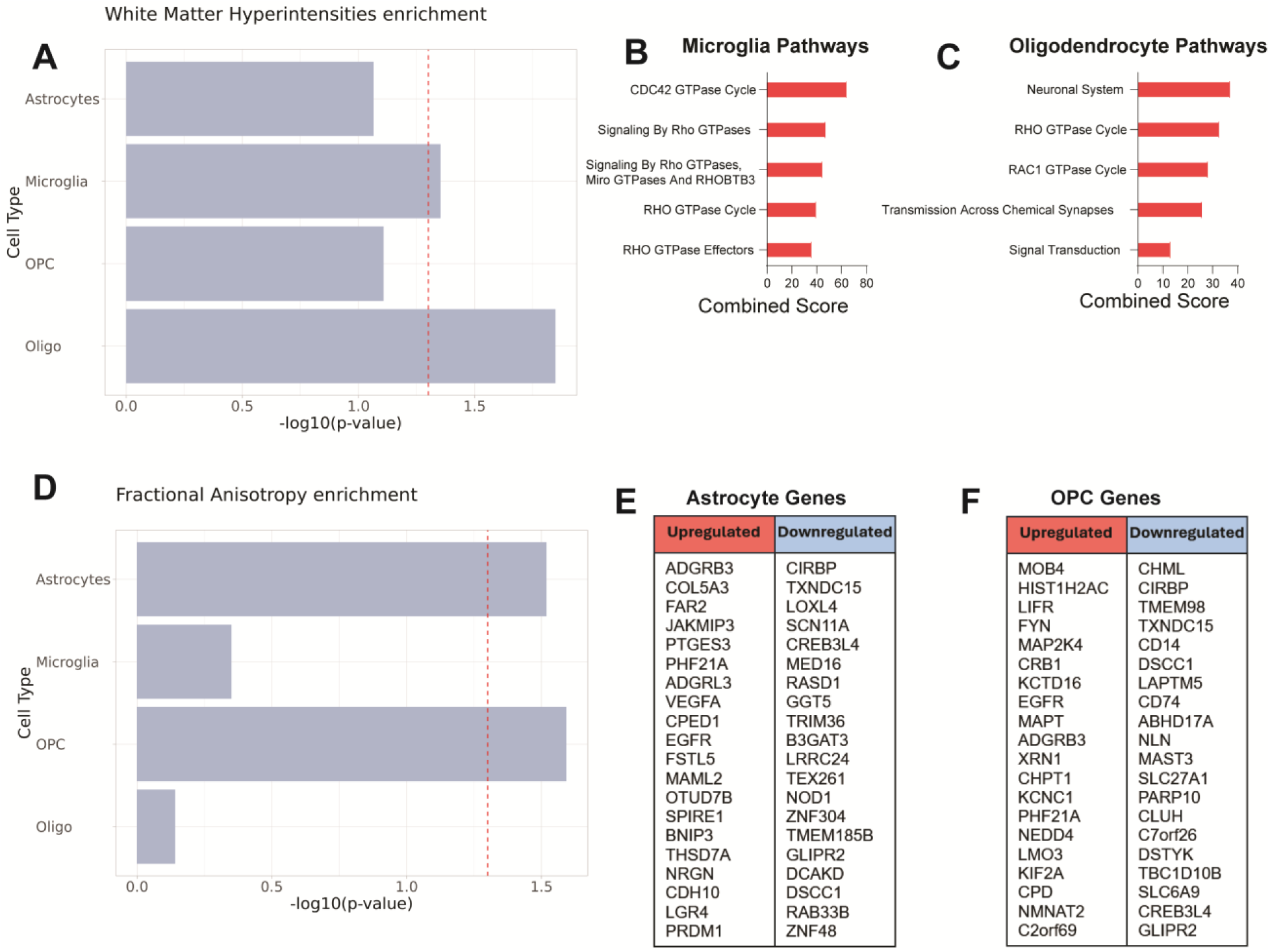
Expression analysis of white matter disease risk variants from GWAS data using MAGMA. (**A**) Association of cell type transcriptomes in VaD with known genetic risk variants linked to increased white matter hyperintensity burden. (**B, C**) Enriched Reactome gene sets obtained from upregulated and downregulated genes in Microglia and Oligodendrocytes linked to increased WMH. (**D**) Association of cell type transcriptomes in VaD with known genetic risk variants linked to decreased global fractional anisotropy. (**E, F**) List of upregulated and downregulated genes in Astrocytes and OPCs linked to decreased global FA (no significant hits found on Reactome).

MAGMA identified microglia and oligodendrocyte transcriptomes as significantly associated with WMH volume (Figure 9A). Looking at the expression of the genes identified by MAGMA, we saw an enrichment of RHO GTPase signaling in both cell types (Figure 9, B and C). Oligodendrocytes also showed increased Transmission Across Chemical Synapses and Signal Transduction. MAGMA also identified astrocyte and OPC transcriptomes as significantly associated with FA (Figure 9D). While several genes from the FA GWAS were found to be differentially expressed across these cell types, no pathways were significantly associated with these gene lists (Figure 9, E and F).

Because of the known overlap between AD pathology and VaD, we extended our analysis to genetic risk variants linked to AD (Figure 10A). Only the oligodendrocyte transcriptome showed significant association with AD genetic risk variants. The genes of interest identified by MAGMA in oligodendrocytes were involved in increased signal transduction, decreased mRNA transcription and decreased DNA repair mechanisms (Figure 10B). We also sought to explore possible association with genetic risk variants associated with intracerebral hemorrhage (ICH), one of the most severe phenotypes of CSVD (Figure 10C). MAGMA identified only the astrocyte transcriptome as being significantly associated with ICH genetic risk variants. While MAGMA identified several genes from the ICH GWAS to be differentially expressed in our astrocytes, no pathways were significantly associated with these gene lists (Figure 10D).

**Figure 10.**
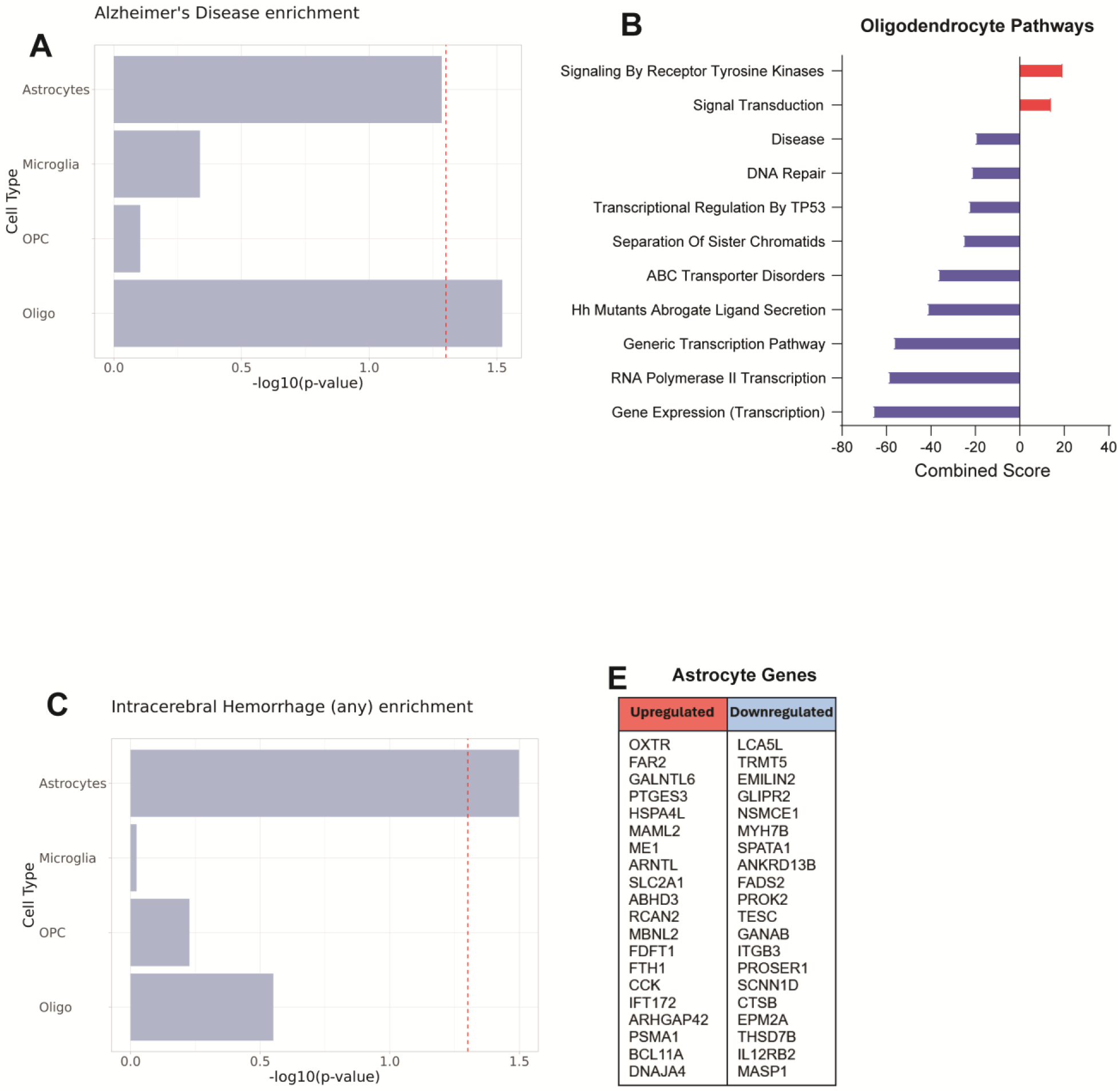
Gene-set analysis of AD and ICH GWAS data using MAGMA. (**A**) Association of cell type transcriptomes in VaD with known genetic risk variants linked to development of Alzheimer’s disease. (**B**) Enriched Reactome gene sets obtained from upregulated and downregulated genes in Oligodendrocytes linked to development of AD. (**D**) Association of cell type transcriptomes in VaD with known genetic risk variants linked to intracerebral hemorrhage. (**E, F**) List of upregulated and downregulated genes in Astrocytes linked to ICH (no significant hits found on Reactome).

## Discussion

Cerebrovascular disease is a driving factor of VaD and has been shown to exacerbate other dementias including Alzheimer’s disease. Thus, there is a pressing need to better understand the mechanisms by which CVD contributes to disease progression in dementias. In our current study, we focus on VaD pathology driven by CSVD leading to periventricular white matter disease. Using snRNAseq of unsorted nuclei isolated from human *postmortem* PVWM, we performed a comprehensive transcriptomic analysis with the goal of identifying biological mechanisms related to CSVD-driven VaD pathology. We compare these findings with those previously reported (25) and then test whether exposure of iMGL to OGD can recapitulate key parts of the VaD microglial transcriptome. Finally, we employ MAGMA to show that our VaD transcriptomic signatures are associated with genetic risk variants identified by GWAS in the white matter CSVD measured by FA and WMH.

Our snRNAseq approach yielded all expected CNS cell populations with an abundance of oligodendrocytes, astrocytes, microglia and OPCs. Similar to previously published results (25), we did not see significant shifts in the proportion of these cells between Control and VaD groups. The total number of predicted receptor-ligand interactions was reduced in VaD, suggesting that an overall cell-cell signaling decrease may be a feature of the lesion site in VaD, possibly as a consequence of chronic regional hypoperfusion. While astrocytes, microglia and oligodendrocytes showed decreased cell-cell signaling in VaD, OPCs showed an increase in signaling. OPCs have been shown to contribute to remyelination by differentiating into mature myelinating oligodendrocytes. Whether OPCs attempt to do this at the white matter lesion sites in VaD remains an open question and will require further testing.

Astrocytes in homeostasis perform multiple supportive functions including the maintenance of the glia limitans (56), ion homeostasis and protection against glutamate-mediated excitotoxicity (57). In response to injury, astrocytes can promote neuroinflammation by producing chemokines (58), releasing reactive oxygen species (59), and disrupting the blood-brain barrier (BBB) via VEGF production (60). For the most part, pathways upregulated in astrocytes in VaD were neither related to their homeostatic functions nor to promoting neuroinflammation. Instead, we observed an increase in pathways related to cellular stress including HSP responses, cell death signaling, cellular senescence and histone-mediated transcriptional repression. Many of these features can be induced by chronic exposure to OGD (61). We also see a decrease of receptor-ligand interactions involved in angiogenesis (ANGPT), efferocytosis (PROS) and anti-inflammatory signaling (MIF), suggesting broad impairment of astrocytic support functions. Finally, we observe a marked upregulation of cadherins (core genes driving the “Reactome Adherens Junction Interactions” pathway R-HSA-418990) as well as receptor-ligand interactions involved in inhibition of axonal growth. In the context of CNS injury, reactive astrocytes engage in fibrotic glial scarring, a process that aims to limit injury spread (62–65). If inflammation is properly resolved, the glial scar can be pivotal to proper wound healing (66, 67), but under chronic conditions it can prevent wound healing by inhibiting axonal growth into the lesion site (68, 69). Overall the transcriptomic signature identified in astrocytes in VaD is reminiscent of reactive astrocytes present in chronic unresolved glial scars (70). Further characterization of chronic ischemic lesions in VaD will be pivotal to understand the role glial scarring may play in disease pathology.

Microglia are the tissue-resident macrophages of the CNS. In homeostasis, microglia perform numerous functions essential for CNS health including rapid surveillance of their environment, efferocytosis, clearance of amyloid and synaptic pruning. In response to injury or infection, microglia quickly change to an ameboid morphology and migrate to the site of injury where they produce inflammatory cytokines and chemokines and perform phagocytosis. In contrast, microglia from subjects with VaD were mainly characterized by a loss of effector functions. They show a global downregulation of translation, transcriptional repression, decreased signals related to amyloid clearance (TREM2, CD74), decreased TAM receptor-mediated efferocytosis (PROS) and decreased production of both pro (LGALS9) and anti-inflammatory (MIF) mediators. They also show increased NMDA receptor-mediated signaling, which may be indicative of excitotoxicity, a well-characterized consequence of chronic ischemia (71).

To test whether OGD exposure could partially recapitulate the microglial transcriptomic signature observed in VaD, we performed single cell RNA sequencing of human iPSC-derived microglia exposed to 24 hrs of OGD. OGD-exposed microglia showed a global decrease in translation and increased transcriptional repression, which partially recapitulates the microglial transcriptome observed in VaD and thus may be one of the earliest responses to hypoperfusion. The iMGL transcriptome in OGD also differed from the microglial transcriptome in VaD in significant ways, with OGD-exposed microglia inducing increased innate immune signaling, increased phagocytic activity, increased ROS production and decreased HSP responses. While significant similarities exist between the microglial response to acute OGD and the microglial transcriptome in VaD, this *in vitro* system does not fully recapitulate the microglial transcriptomic signature in VaD. Given the partial overlap found with iPSC-derived microglia, further studies using mixed glial and neurovascular unit cultures and brain organoids may be exciting avenues to explore for improved *in vitro* modeling of disease.

Proper myelination of neuronal axons by oligodendrocytes in the white matter is essential for cognitive function (72). Increased demyelination has been observed in both AD and VaD and may play a key role in VaD-driven cognitive dysfunction (73). As expected, receptor-ligand interactions implicated in myelination such as MAG, CLDN11 and NRG were downregulated in oligodendrocytes from subjects with VaD. We also observe a dramatic decrease in oxidative phosphorylation, decreased protein trafficking and increased transcriptional repression which may reflect OGD-driven stress in these cells. Human mature oligodendrocytes have been shown to be surprisingly resistant to metabolic stress, in part due to their ability to decrease their metabolic rate under these conditions (74).

Somewhat surprisingly, oligodendrocytes in VaD also show an upregulation in cholesterol biosynthesis when compared to controls. This is somewhat paradoxical since anabolic mechanisms, such as cholesterol biosynthesis, are usually downregulated in response to metabolic stress (75). Cholesterol is essential for myelination and synapse formation (76, 77), and the inability of lipoproteins to cross the blood-brain barrier means that maintenance of brain cholesterol levels largely depends on parenchymal cholesterol biosynthesis (78). In acute ischemia, reactive astrocytes have been shown to increase cholesterol biosynthesis to supply myelinating oligodendrocytes (79, 80). We do not observe increased cholesterol biosynthesis in astrocytes in VaD, but we do observe it in microglia and oligodendrocytes in VaD. This raises the possibility that oligodendrocyte-mediated cholesterol biosynthesis may be a stress resistance mechanism under chronic conditions to support oligodendrocyte survival in the absence of astrocyte-derived cholesterol. Taken together these results suggest an oligodendrocyte transcriptome in VaD defined by increased cellular stress, decreased myelination and increased cholesterol production, suggesting a shift towards prioritizing cell survival over cell function.

As previously stated, OPCs were the only cell type that showed a global upregulation of cell-cell signaling in VaD. Unlike mature oligodendrocytes, OPCs are motile, and are well-known for their involvement in remyelination of white matter lesions (77). This raises the possibility that OPCs captured in VaD samples have not been chronically exposed to ischemia and are instead migrating from non-affected regions of the brain in an attempt to remyelinate the lesion site. OPCs in VaD show features related to cell stress including an increase in HSP responses and NMDA receptor signaling. However, OPCs also show increased FGF receptor-ligand interactions primarily those mediated by FGF2, which has been found to protect oligodendrocyte precursors from OGD *in vitro* (81). FGF signaling is also known to promote OPC-mediated remyelination (42, 81). Taken together these results suggest that OPCs in VaD may be migrating to the lesion site in an attempt to remyelinate but upregulating several cellular stress responses as they are exposed to chronic ischemia.

Multiple factors outside of CSVD may contribute to the transcriptomic profiles we characterized in VaD. To try to determine whether CSVD was significantly associated with our transcriptomic profiles in VaD, we tested whether genetic risk variants predictive of CSVD were significantly associated with any of our cell type-specific transcriptomic profiles in VaD. Structural deterioration of the white matter can be measured via MRI, in the form of white matter hyperintensities (T2-weighted) or fractional anisotropy (diffusion-weighted) (52). Using MAGMA, we saw that differential gene expression profiles in microglia and oligodendrocytes were associated with increased WMH burden, while differential gene expression profiles in astrocytes and OPCs were associated with decreased FA, reflecting compromised white matter microstructural integrity. In contrast, genetic risk variants from small vessel stroke and enlarged perivascular spaces, phenotypes that are associated to cognitive decline but are not white matter specific, did not show significant association with any of our cell type-specific VaD transcriptomes. These results suggest PVWM disease may have unique transcriptomic profiles in subcortical lacunar infarctions in VaD. This would be consistent with models of PVWM disease involving unique mechanisms of breakdown of the ependymal lining of the ventricles and dysregulated fluid flow from the ventricles (23, 82), In contrast, deep subcortical white matter lesions are more closely associated with hypoxia, microglial activation, arteriosclerosis and enlarged perivascular spaces (23, 83, 84). Additional analyses using GWAS for AD and ICH showed some association with the transcriptomic profiles characterized in VaD, suggesting there may be some overlap in the molecular mechanisms driving PVWM disease and those driving AD and ICH.

The transcriptomic profiles have characterized in VaD are marked by an overall increase in cellular stress responses and impairment of cell type-specific functions pivotal to CNS homeostasis. Using a previously published snRNAseq study, we tested whether these cell type-specific transcriptomic signatures were conserved across datasets. RRHO analysis showed the upregulation of HSP responses in VaD across both datasets. This was confirmed via module scoring which showed increased HSP response across datasets but no increase in the UPR module in either dataset. While the HSP response and UPR are both mechanisms that deal with proteotoxic stress in the cell, they do so in different cell compartments with the HSP response targeting the cytosol and the UPR targeting the ER (85). Increased HSPs in response to ischemia and it’s cytoprotective effects have been well documented (86–91). Interestingly, one of the ways in which HSP response mediates cytoprotection is by inhibiting several steps of the apoptotic signaling pathway (92–97). Network analysis shows a common upregulation of cellular senescence and apoptotic signaling in astrocytes in VaD across both datasets. Overall, this overlap across two independent datasets strongly suggests that increased cellular stress is a conserved feature of the cellular transcriptome in PVWM lesions in VaD and that increased HSP response may be a resilience mechanism employed by cells at the lesion site to protect against chronic ischemia.

Despite a common overlap across datasets, there remains significant differences between them, particularly in regard to neuroinflammatory signaling and protein translation. Mitroi et al observed an increase in both neuroinflammatory signaling and protein translation in VaD while we observed the opposite. To ensure these differences were not due to the data analysis pipeline used, we reprocessed the raw data from the *Mitroi et al* dataset using our pipeline and were able to recapitulate the general UMAP structure, identify the same cell types and found broadly similar results to those in the original publication (25). One difference between the protocols used is that we sequenced all isolated nuclei after sucrose ultracentrifugation while the Mitroi et al team sorted their isolated nuclei based on DAPI (or DAPI + ERG) expression prior to sequencing. While this difference is worth noting, both teams used similar nuclei isolation protocols and sampled white matter from the same brain regions. This suggests that there may be meaningful biological differences between the VaD samples processed by both teams which, given the heterogeneity of disease, is not entirely surprising.

Further insights into the pathology of VaD warrants substantial future research into the different factors that converge to promote VaD pathology, further characterization of disease progression and validation of key biological mechanisms identified via transcriptomic studies of VaD. Our transcriptomic findings partially align and partially differ from prior research. We observe a marked upregulation of genes related to the HSP response across all cell types in VaD, which aligns with previous transcriptomic findings (25) and raises the possibility of the HSP response being a promising therapeutic target. Unlike previous studies, we do not observe upregulated neuroinflammatory pathways in VaD, nor did we identify infiltrating leukocytes. Many previous studies have strongly suggested increased neuroinflammation may play a role in VaD. While we did not observe upregulated neuroinflammation at the lesion site in our VaD cohort, we submit that this discrepancy may be a result of our dataset providing a ‘late snapshot’ of disease pathology at autopsy. Inflammatory responses require substantial energetic resources which, under chronic ischemic conditions may become unsustainable over time, causing a shift of the cellular landscape from trying to resolve injury to prioritizing cell survival. Finally, we did not enrich for vascular cells and did not identify pericytes, vascular smooth muscle cells, or endothelial cells. These components of the neurovascular unit warrant further study, particularly at early stages of CSVD.

Overall, we contribute a transcriptomic dataset as a resource for understanding VaD. We report novel findings in VaD involving the marked upregulation of several pathways related to cell stress such as: cell death signaling, cellular senescence, decreased oxidative phosphorylation, transcriptional repression and decreases in protein translation. We show significant association of CSVD genetic risk variants to our transcriptomic signatures in VaD. We show that iMGL exposure to 24 hrs of OGD partially recapitulates the microglial transcriptomic signature observed in VaD suggesting OGD may be a driving factor of VaD pathology. Finally, we leverage a previously published snRNAseq dataset of VaD-affected postmortem PVWM and detail a shared signature despite disease heterogeneity which could represent a promising therapeutic target.

## Methods

### Sex as a biological variable

Our study design considered both male and female sex. Human postmortem samples are balanced for sex and primary human iPSCs cultures include donor cells from one male and one female donor.

### Sample Collection and Single-nucleus RNA sequencing

Our dataset contains data from 8 post-mortem samples of human PVWM. Samples were acquired through the NIH NeuroBioBank (Request ID #1458). Selected samples were matched for age and sex and limited to those without neurofibrillary tangles or Braak & Braak stages I or II (Supplementary Table 1).

Nuclei extraction was performed as previously described (98) and all steps were performed at 4°C. In brief, 50 mg of frozen PVWM were mixed with 15 mLs of homogenization buffer and dounce homogenized 10-15 times using the loose pestle and 10-15 times using the tight pestle (Wheaton Cat # 357544). The tissue suspension was then subjected to sucrose ultracentrifugation (Beckman Coulter Avanti JXN-26) at 24,000 RPM for 1 hr. Supernatant was discarded and nuclei were resuspended in chilled 1X PBS at a concentration of 1000 nuclei/uL. Nuclei were then processed using Chromium Single Cell 3’ v3.1 technology (10X Genomics) and sequenced using a NovaSeq 6000 sequencer.

### Alignment and Pre-Processing

Raw sequencing data was aligned to the human genome hg38 using Cell Ranger v7.0.0 with include-introns set to TRUE. Data was processed using Seurat v5 (99, 100).

Several filters were used to exclude doublets and dying cells. Nuclei with unique feature counts over 6,000 or below 500 were removed as well as nuclei that contained higher than 5% mitochondrial counts. Data was scaled and normalized using SCTransform v2 and percentage of mitochondrial counts was regressed out using the vars.to.regress parameter. The first 50 principal components were calculated and used to integrate samples using the CCAIntegration method. Clustering was done using the Louvain algorithm and we generated UMAPs based on the first 40 principal components at a resolution of 0.1.

### Cluster Annotation and identification of Differentially Expressed Genes

Clusters were manually annotated based on their expression of previously validated cell type-specific markers (101). Counts were pseudobulked by sample and cell type using the aggregate.Matrix function. Two filters were added to the function filterByExpr: min.count = 0.1 and min.total.count = 15. DEGs were then determined using the EdgeR’s voomLmFit approach (102).

### Gene Set Enrichment Analysis

Pseudobulk DEGs were sorted by their F-statistic value and submitted for analysis using the GSEA v4.3.2 software (40). We used gene sets from Reactome via MSigDB (v2023.2.Hs) and excluded those larger than 500 genes or smaller than 15 genes.

Analysis was performed using 1000 permutations and unless otherwise stated, results were filtered by an FDR < 0.05.

### Cytoscape Network Analysis

Network analysis was performed using Cytoscape v3.10.2 software and three plug-ins: EnrichmentMap (v3.4.0), clustermaker2 (v2.3.4) and AutoAnnotate (v1.5.0). GSEA results were uploaded onto EnrichmentMap for visualization. Each pathway appears as a ‘node’ while pathways sharing common genes are connected to each other via ‘edges’. Using the Markov Clustering (MCL) algorithm, clustermaker2 grouped related GSEA pathways together into annotations based on their similarity coefficient. Finally, we used AutoAnnotate to create placeholder labels based on the most common keywords shared among the pathway descriptions provided by Reactome. While helpful, these names were not always accurate. We manually checked the member pathways of each cluster and the genes driving them to ensure cluster names reflected the biological theme of the annotated cluster. When appropriate, these clusters were renamed.

### Predicted Receptor-Ligand Interaction Analysis using CellChatDB

We performed predicted receptor-ligand interaction analysis using the CellChat v2 R package and the human CellChatDB database which contains over 3,000 validated interactions between ligands, receptors and cofactors (103). We limited this analysis to four distinct celltypes: astrocytes, microglia, oligodendrocytes and OPCs. Using our pre-processed Seurat objects as input, CellChat inferred cell-cell communication using cell type-specific expression data of receptor-ligand genes and assigned each interaction a communication probability value.

Using the rankNet function, we compared the interaction strength of each signaling pathway across conditions. Setting the do.stat parameter to TRUE, a paired Wilcoxon test was performed to determine whether the difference in information flow between both conditions is significant. Only significant interactions with a p-value < 0.05 and with consistent receptor-ligand expression observed by VlnPlots are shown in our final figures. From the signaling pathways that met these criteria, we selected specific receptor-ligand interactions of interest and visualized their communication probability via Bubble plots using the netVisual_Bubble function.

### PMI Correlation Analysis

To assess the extent to which post-mortem interval may have driven the cellular response to heat stress, we selected the core enriched genes from three Reactome pathways related to this function: HSF1 activation (R-HSA-3371511), HSF1 dependent transactivation (R-HSA-3371571), and Attenuation Phase (R-HSA-3371568). We tested the correlation of each gene’s expression by sample with the PMI values using Pearson’s Correlation Coefficient test.

### In vitro culture of human iPSC-derived microglia (iMGL)

Human induced pluripotent stem cells (iPSCs) were cultured in mTeSR Plus (STEMCELL Technologies, Cat# 100-0276) on Geltrex-coated (Gibco, Cat# A1569601) 6-well plates. Colonies were dissociated using ReLeSR (STEMCELL Technologies, Cat# 05872) into smaller aggregates, with approximately 50 aggregates plated per well in a 6-well plate. Aggregates were cultured in STEMdiff Medium A (STEMCELL Technologies, Cat# 05310) from day 0 to day 2 and in STEMdiff Medium B (STEMCELL Technologies, Cat# 05310) from day 3 to day 12 to differentiate them into hematopoietic progenitor cells (HPCs). Floating HPCs were collected from the medium on day 12 and plated at a density of 150,000 to 200,000 cells per well in STEMdiff Microglia Differentiation Medium (STEMCELL Technologies, Cat# 100-0019) to induce microglial differentiation. The medium was topped with 1 mL every other day, and cells were subcultured every 8 days. On day 24 of incubation with STEMdiff Microglial Differentiation Medium, cells were transferred to home-brewed Microglia Maturation Medium for 5 days before treatment.

In brief, maturation media consisted of DMEM/F-12 (Elabscience, Cat# PM150322) supplemented with 1x ITS-G (Gibco, Cat# 41400045), 200 µM Monothioglycerol (MTG) (Sigma, Cat# M6145), 4 µg/mL Insulin (Sigma, Cat# I9278), 100 ng/mL Interleukin-34 (IL-34) (Peprotech, Cat# 200-34), 50 ng/mL Transforming Growth Factor Beta (TGF-β) (Peprotech, Cat# 100-21), 25 ng/mL Macrophage Colony-Stimulating Factor (M-CSF) (Peprotech, Cat# 300-25), 100 ng/mL Fractalkine (CX3CL1) (Peprotech, Cat# 300-31), and 100 ng/mL CD200 (R&D Systems, Cat# 2724-CD050). On treatment day, maturation media for the oxygen-glucose deprivation (OGD) treatment was made using glucose-free DMEM/F-12 (Elabscience, Cat# PM150312). Oxygen was set to 1% for hypoxia and OGD treatments using a hypoxic chamber while normoxic controls were incubated under default conditions. After 24 hrs of exposure to treatment, cells were gently washed and resuspended in chilled 1X PBS at a concentration of 1000 cells/uL. Cells were then processed using Chromium Single Cell 3’ v3.1 technology (10X Genomics) and sequenced using a NovaSeq X Plus sequencer.

### Data analysis of iMGL single-cell RNA sequencing

Aside from two main differences detailed here, data analysis for scRNAseq was the same as that for snRNAseq. During pre-processing, cells with unique feature counts over 9,000 or below 1,500 were removed as well as cells that contained higher than 15% mitochondrial counts. Instead of pseudobulking, DEGs were determined using Seurat v5’s Wilcoxon Rank Sum test implemented with the FindMarkers() function.

### Canonical Pathway Analysis and Comparison Analysis via IPA

Lists of differentially expressed genes by cell type were used as input for Ingenuity Pathway Analysis (IPA). Gene lists were filtered for adjusted p-value ≤ 0.05 and Canonical pathway analysis was performed using standard analysis settings. Results were excluded if driven by 5 or less genes or if −log(p-value) was below 1.30. For IPA comparison analysis, we only included canonical pathways with an absolute value z-score ≥ 1 in both datasets.

### Testing overlap of snRNAseq datasets using Rank-Rank Hypergeometric Test (RRHO)

Pre-ranked gene lists sorted by F-statistic were generated as previously described for astrocytes, microglia, oligodendrocytes and OPCs from our dataset and the *Mitroi et al* dataset. Overlap for each cell type in both datasets was determined using the RRHO function’s two-sided test from the RRHO v1.42 R package. Results are shown as a heatmap with a signed log-transformed hypergeometric p-value scale. Upregulated and downregulated genes identified as overlapping by RRHO were submitted to Enrichr (104) and cross-referenced to the Reactome database. Matching hits were sorted by adjusted p-value ≤ 0.05 and removed if they were driven by less than 8 total genes. Top 10 hits were sorted by the signed Combined Score provided by Enrichr.

Since the iMGL dataset consisted of one 10X run, this data was not pseudobulked. To assess overlap between iMGL, our dataset from VaD-affected subjects, and *Mitroi et al* microglia, pre-ranked lists were sorted by the fold change value obtained from Seurat’s Wilcoxon Rank Sum test.

### Generating module scores across datasets

Gene sets from MSigDB (v2023.2.Hs) were used as modules to perform module scoring. We used Seurat v5’s AddModuleScore function which determines the average expression of a given module at the single-cell level (47). Module scoring was done on our snRNAseq dataset as well as the DAPI-sorted and DAPI+ERG-sorted *Mitroi et al* datasets.

### Determining the association of cell type transcriptomes in VaD with known genetic risk variants using MAGMA

To determine whether genetic risk variants associated to CSVD were overexpressed in any of our cell type-specific VaD transcriptomic signatures, we used Multi-marker Analysis of GenoMic Annotation (MAGMA, v.1.10) (48). Differentially expressed genes by cell type were generated as previously described using the EdgeR package. For each GWAS, we first mapped all genetic variants to genes based on the publicly available gene reference file provided by MAGMA (https://ctg.cncr.nl/software/magma, accessed on 22 August 2024), which includes the genomic location of 19,427 genes. We then performed a gene analysis using the SNP-wise Mean model to calculate an aggregated p-value for each gene. The 1000 Genomes European panel was employed as a reference to assess the linkage disequilibrium between SNPs. Finally, we conducted MAGMA’s gene property analysis, which is similar to a competitive gene-set analysis but uses a continuous variable as the predictor rather than a binary gene set. The continuous variable used in our analysis was the logarithm of the differential expression p-value obtained from EdgeR DEG analysis, signed by the direction of the log fold change.

For cell types whose transcriptomes were found to have significantly enriched GWAS-identified genes, we took the upregulated and downregulated genes with p-values ≤ 0.05, submitted them to Enrichr (104) and cross-referenced to the Reactome database. Matching hits were sorted by adjusted p-value ≤ 0.05 and removed if they were driven by less than 8 total genes. Top hits were sorted by the signed Combined Score provided by Enrichr. For cell types that did not have significant Enrichr hits, we show a table with the top 20 most upregulated and downregulated genes.

### Statistics

Graphs were created using the ggplot in R or GraphPad Prism 9. To determine differentially expressed genes we used either EdgeR’s voomLmFit approach (102) or Seurat’s Wilcoxon Rank Sum test approach (99, 100). For PMI correlation analysis we used Pearson’s Correlation Coefficient test. Adjusted p-values were obtained using the Bonferroni correction for multiple comparisons.

### Study approval

Our study was approved by the Yale IRB (1506016023).

### Data availability

Raw sequencing data is accessible via the National Center for Biotechnology Information Gene Expression Omnibus (GSE282111 and GSE282112). R Scripts used to generate figures can be found on GitHub (SansingLab/SDP).

## Author Contributions

Designed project SDP, MHA, and LHS.

Designed experiments SDP, CL, MHA, LZ, and LHS.

Conducted experiments SDP, CL, and LZ.

Processed data SDP, JHD, CAR, CL and BZ.

Interpreted Results SDP, JHD, CAR, BZ, LZ, KJB, and AJM.

Wrote manuscript SDP, CAR and CL.

Edited manuscript SDP, JHD, CAR, CL, AJM and LHS.

Procured funding SDP, KJB and LHS.

## Acknowledgements

This work was supported by NIAID T32 AI007019 (SDP), American Heart Association Research Supplement to Promote Diversity in Science #000293492054 (SDP), AAN/AHA Ralph L. Sacco Scholars Fellowship 24RSSPOST1328228 (CAR), R01AG068030, RF1AG065926-01A1, R56AG071291, R01ES033630 (KJB), American Heart Association 19EIA34770133 (LHS). Sequencing was supported by the NIGMS/NIH 1S10OD030363-01A1.

We thank the NIH NeuroBioBank at Harvard University, the University of Miami and the Icahn School of Medicine at Mount Sinai for collecting and making available the human tissue samples used in this publication and the patients and families who made the donations. We also thank Rolando Garcia Milian, Prashant Emani, and Ran Meng for advice on bioinformatics analyses.

---

**Figure S1.**
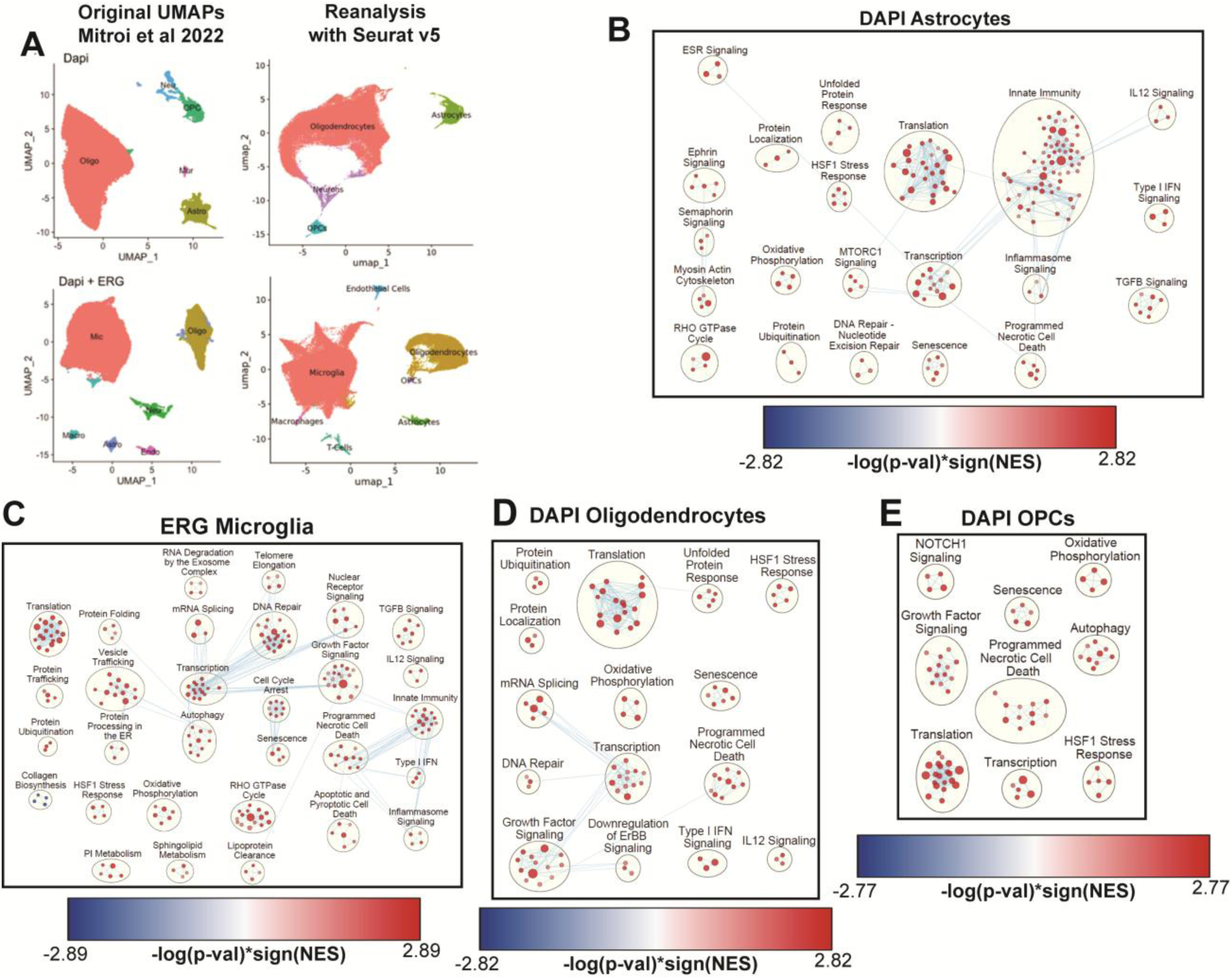
GSEA Analysis of Mitroi et al VaD datasets. (**A**) Side by side comparison of Mitroi et al UMAPs from original publication (left) and after reanalysis with our Seurat v5 pipeline (right). (**B-D**) Cytoscape EnrichmentMap showing individual GSEA hits from Reactome (FDR < 0.05) clustered via AutoAnnotate.

**Figure S2.**
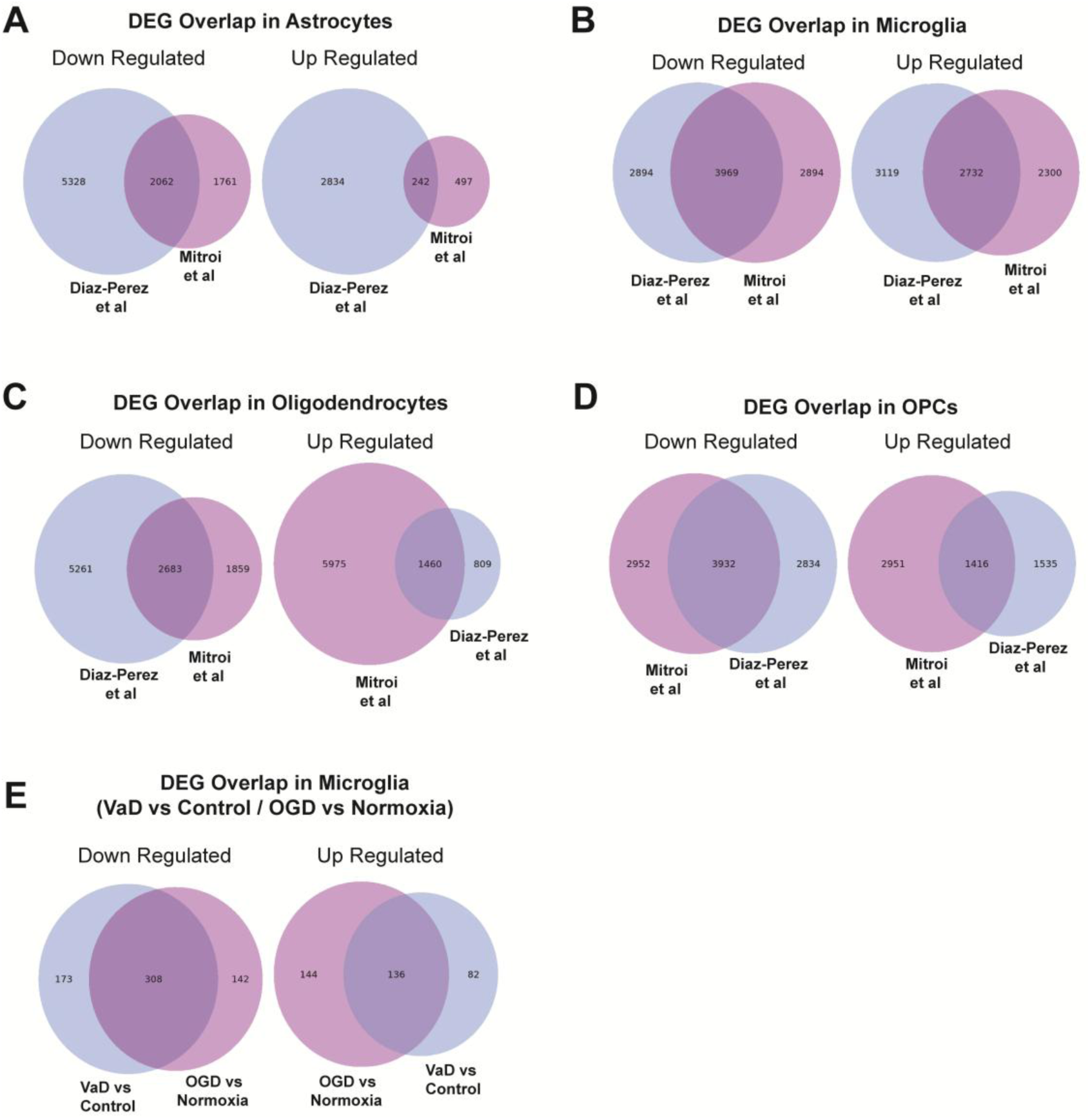
Overlapping genes identified by RRHO in comparisons between transcriptomic datasets. Venn diagrams showing overlap of pre-ranked gene lists (VaD vs Control) from (**A**) astrocytes, (**B**) microglia, (**C**) oligodendrocytes and (**D**) OPCs from out dataset (*Diaz-Perez et al*) and the *Mitroi et al* dataset. Venn diagram showing overlap of preranked gene lists between microglia from VaD vs Control and OGD vs Normoxia.

**Figure S3.**
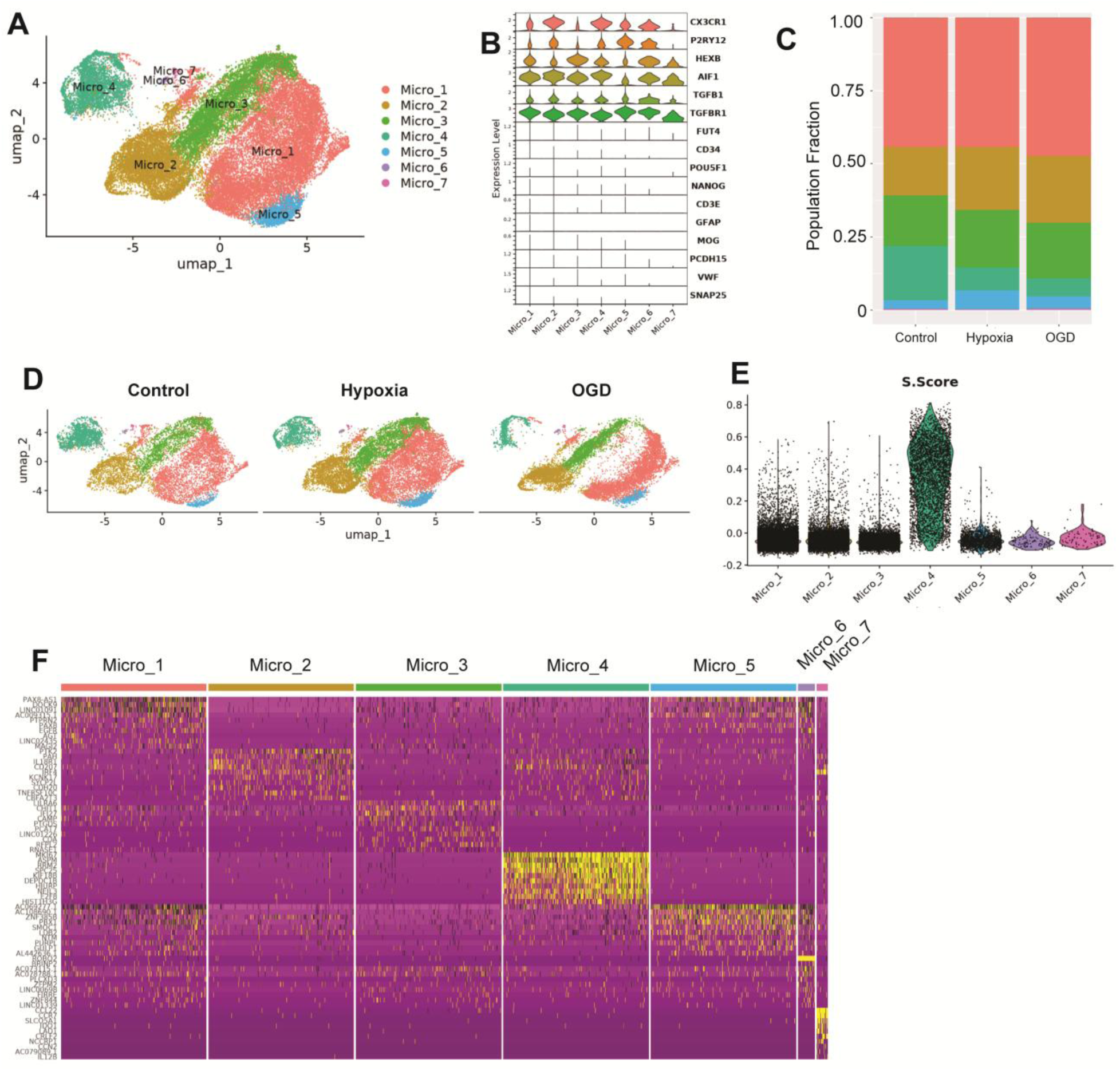
Single cell landscape of iMGL. (**A**) UMAP of iMGL dataset. (**B**) Violin plot of multiple microglia-specific, iPSC-specific and other CNS cell type-specific markers in iMGL dataset. (**C**) Cluster proportions split by normoxia, hypoxia and OGD. (**D**) UMAP of iMGL dataset split by normoxia, hypoxia and OGD. (**E**) Module scoring of each cluster using cell cycle genes. (**F**) Heatmap of top 10 markers that distinguish each cluster from all other cells in our dataset.

**Figure S4.**
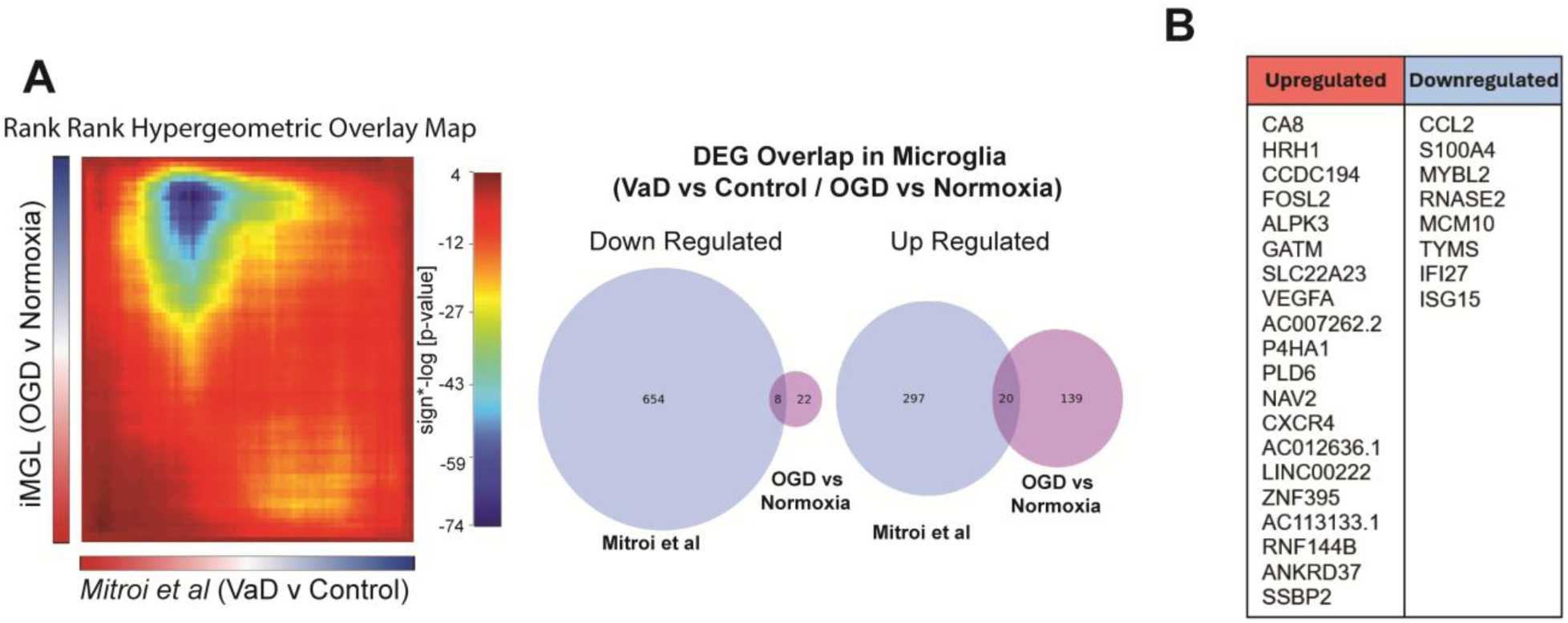
Anticorrelation of OGD-exposed iMGL and *Mitroi et al* dataset. (**A**) RRHO heatmap and Venn diagrams showing overlap of pre-ranked gene lists from *Mitroi et al* (VaD vs Control) and iMGL (OGD vs Normoxia) datasets. Negative values on the −log(p-val) scale is indicative of an almost perfect anti-correlation between the lists of differentially expressed genes from *Mitroi et al* and iMGL. (**B**) List of overlapping upregulated and downregulated genes between *Mitroi et al* and iMGL (no significant hits found on Reactome).

**Figure S5.**
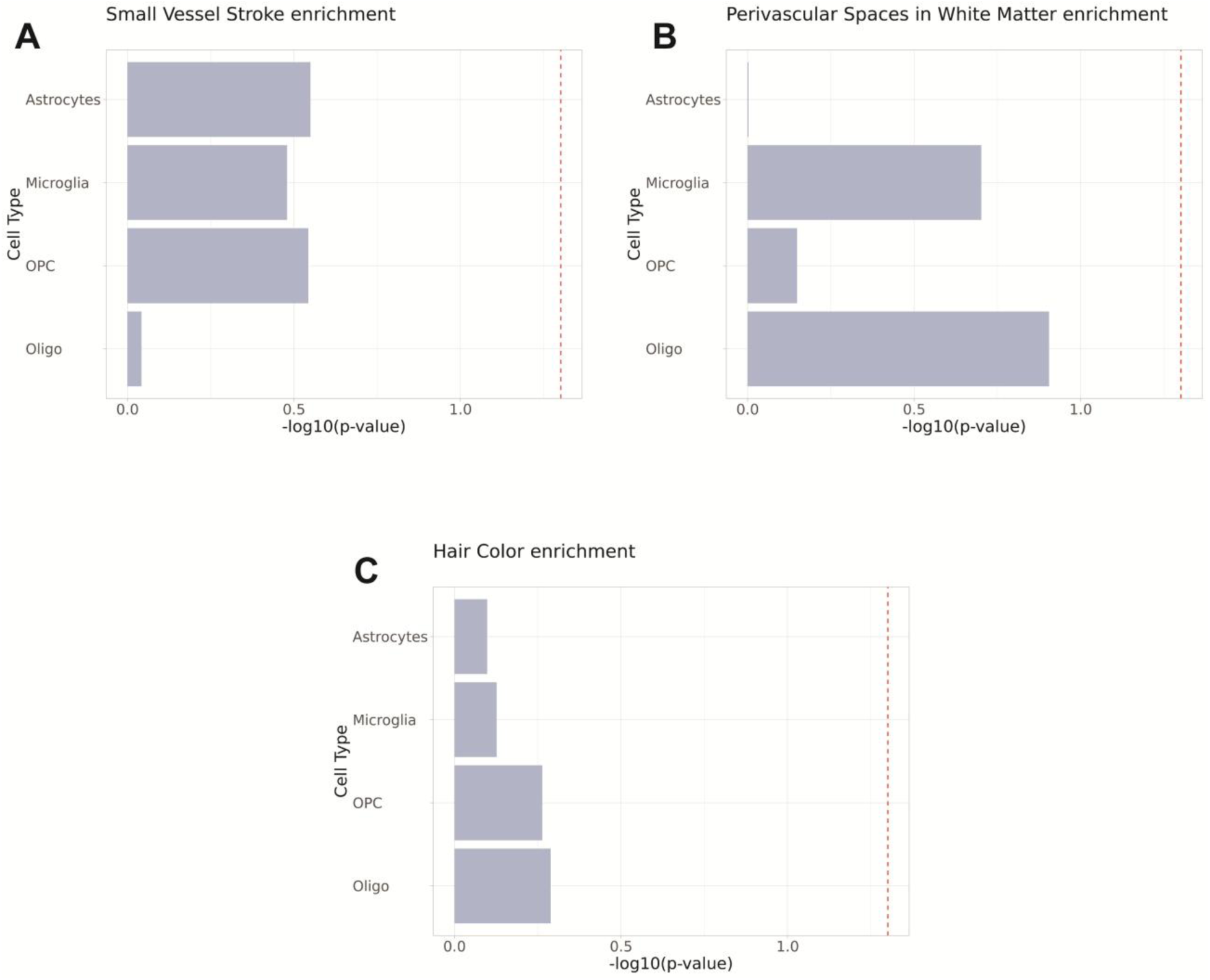
Additional gene-set analysis of GWAS data using MAGMA. Association of cell type transcriptomes in VaD with known genetic risk variants linked to (A) small vessel stroke, (B) enlarged perivascular space and (C) hair color (negative control). The red lines represent the p-value threshold of 0.05, indicating statistically significant associations.

**Supplemental Table 1.**
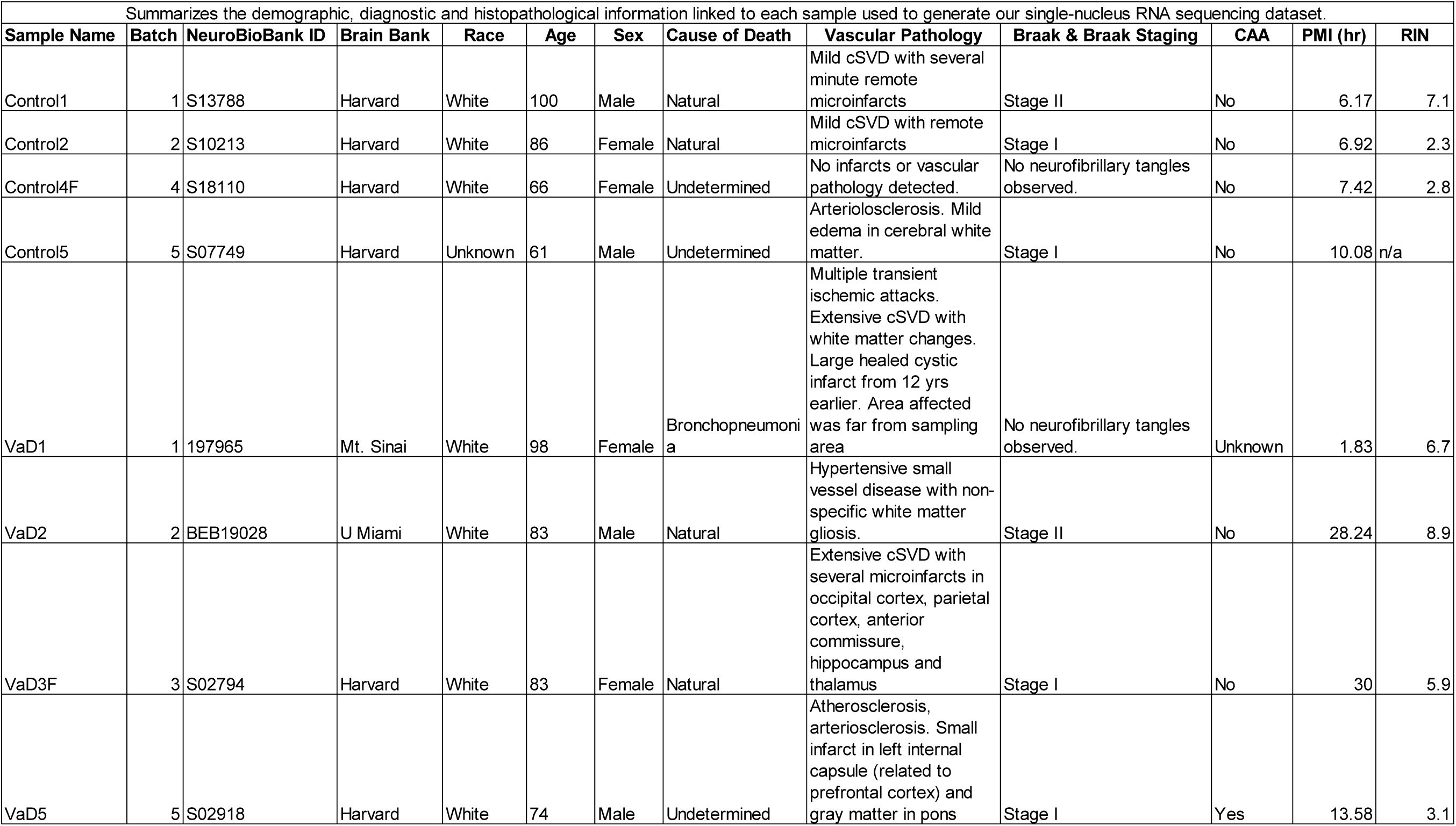
Demographic and histopathological information of periventricular white matter samples.

**Supplemental Table 2.** Quality Control metrics of sequenced snRNAseq samples.

|  |  |  |  |  |  |  |  |  |  |  |  |
| --- | --- | --- | --- | --- | --- | --- | --- | --- | --- | --- | --- |
| Summarizes the quality control parameters obtained from each sample post-sequencing, after CellRanger alignment and after implementing initial QC cutoffs in Seurat v5. |  |  |  |  |  |  |  |  |  |  |  |
| Pre-Sequencing |  |  | Post-Sequencing Alignment |  |  |  |  | Dataset Integration |  |  |  |
| Name | Ave bp | cdNA (ng/uL) | # of Nuclei | Median Genes per Cell | Total Genes Detected | % Mapped to Transcriptome | Average of MT % | Range of MT % | Pre-Filtering | Post-Filtering | % Filtered |
| Control1 | 457 | 1.48 | 5,143 | 2,875 | 29,657 | 69.90% | 0.80% | 0% - 23.09% | 5143 | 4937 | 4.01% |
| Control2 | 515 | 8.61 | 3,320 | 1,826 | 27,139 | 62.20% | 0.28% | 0% - 23.11% | 3320 | 3260 | 1.81% |
| Control4F | 442 | 8.61 | 6,691 | 2,931 | 30,872 | 70.70% | 1.33% | 0% - 48.44% | 6691 | 6420 | 4.05% |
| Control5 | 469 | 5.07 | 2,023 | 2,159 | 26,029 | 36.50% | 0.79% | 0% - 10.06% | 2023 | 1842 | 8.95% |
| VaD1 | 460 | 2.22 | 5,795 | 2,894 | 30,179 | 72.10% | 1.73% | 0% - 56.07% | 5795 | 5476 | 5.50% |
| VaD2 | 493 | 22.5 | 4,784 | 2,288 | 29,355 | 78.00% | 0.38% | 0% - 35.46% | 4784 | 4374 | 8.57% |
| VaD3F | 464 | 3.14 | 3,287 | 2,556 | 30,601 | 67.30% | 1.58% | 0% - 74.16% | 3287 | 2670 | 18.77% |
| VaD5 | 469 | 4.69 | 5,313 | 2,457 | 28,741 | 63.00% | 0.57% | 0% - 39.09% | 5310 | 5177 | 2.50% |

**Supplemental Table 3.** Total number of nuclei contributed by sample.

| **Supplemental Table 3. Total number of nuclei contributed by sample** |  |  |  |  |  |  |  |  |
| --- | --- | --- | --- | --- | --- | --- | --- | --- |
| Shows the count of nuclei obtained from each cluster by sample of origin after quality control filtering. |  |  |  |  |  |  |  |  |
| **Cluster Annotation** | **Control1** | **Control2** | **Control4** | **Control5** | **Vad1** | **Vad2** | **Vad3** | **Vad5** |
| **Oligo_1** | 2511 | 1835 | 2677 | 756 | 3260 | 2283 | 890 | 2522 |
| **Oligo_2** | 1546 | 1024 | 2010 | 477 | 1344 | 1361 | 591 | 1392 |
| **Astrocytes** | 162 | 115 | 771 | 170 | 390 | 250 | 292 | 482 |
| **Microglia** | 364 | 64 | 543 | 60 | 203 | 225 | 155 | 341 |
| **OPCs** | 107 | 157 | 222 | 173 | 129 | 106 | 231 | 330 |
| **Neurons** | 161 | 33 | 135 | 164 | 40 | 122 | 423 | 44 |
| **Oligo_3** | 79 | 21 | 30 | 40 | 97 | 24 | 33 | 41 |
| **Endothelial Cells** | 7 | 11 | 32 | 2 | 13 | 3 | 55 | 25 |

**Supplemental Table 4.**
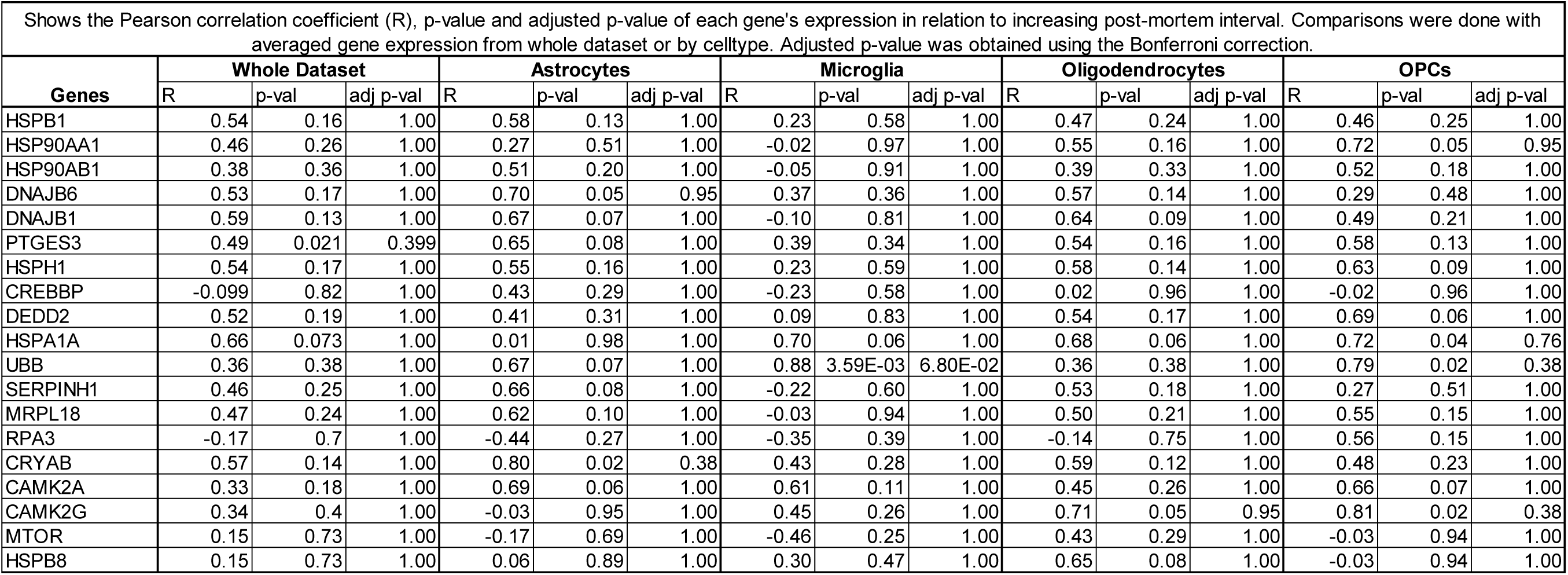
Correlation of heat shock protein gene expression with post-mortem interval.

| Shows the Pearson correlation coefficient (R), p-value and adjusted p-value of each gene's expression in relation to increasing post-mortem interval. Comparisons were done with averaged gene expression from whole dataset or by celltype. Adjusted p-value was obtained using the Bonferroni correction. |  |  |  |  |  |  |  |  |  |  |  |  |  |  |  |
| --- | --- | --- | --- | --- | --- | --- | --- | --- | --- | --- | --- | --- | --- | --- | --- |
| Genes | Whole Dataset |  |  | Astrocytes |  |  | Microglia |  |  | Oligodendrocytes |  |  | OPCs |  |  |
|  | R | p-val | adj p-val | R | p-val | adj p-val | R | p-val | adj p-val | R | p-val | adj p-val | R | p-val | adj p-val |
| HSPB1 | 0.54 | 0.16 | 1.00 | 0.58 | 0.13 | 1.00 | 0.23 | 0.58 | 1.00 | 0.47 | 0.24 | 1.00 | 0.46 | 0.25 | 1.00 |
| HSP90AA1 | 0.46 | 0.26 | 1.00 | 0.27 | 0.51 | 1.00 | -0.02 | 0.97 | 1.00 | 0.55 | 0.16 | 1.00 | 0.72 | 0.05 | 0.95 |
| HSP90AB1 | 0.38 | 0.36 | 1.00 | 0.51 | 0.20 | 1.00 | -0.05 | 0.91 | 1.00 | 0.39 | 0.33 | 1.00 | 0.52 | 0.18 | 1.00 |
| DNAJB6 | 0.53 | 0.17 | 1.00 | 0.70 | 0.05 | 0.95 | 0.37 | 0.36 | 1.00 | 0.57 | 0.14 | 1.00 | 0.29 | 0.48 | 1.00 |
| DNAJB1 | 0.59 | 0.13 | 1.00 | 0.67 | 0.07 | 1.00 | -0.10 | 0.81 | 1.00 | 0.64 | 0.09 | 1.00 | 0.49 | 0.21 | 1.00 |
| PTGES3 | 0.49 | 0.021 | 0.399 | 0.65 | 0.08 | 1.00 | 0.39 | 0.34 | 1.00 | 0.54 | 0.16 | 1.00 | 0.58 | 0.13 | 1.00 |
| HSPH1 | 0.54 | 0.17 | 1.00 | 0.55 | 0.16 | 1.00 | 0.23 | 0.59 | 1.00 | 0.58 | 0.14 | 1.00 | 0.63 | 0.09 | 1.00 |
| CREBBP | -0.099 | 0.82 | 1.00 | 0.43 | 0.29 | 1.00 | -0.23 | 0.58 | 1.00 | 0.02 | 0.96 | 1.00 | -0.02 | 0.96 | 1.00 |
| DEDD2 | 0.52 | 0.19 | 1.00 | 0.41 | 0.31 | 1.00 | 0.09 | 0.83 | 1.00 | 0.54 | 0.17 | 1.00 | 0.69 | 0.06 | 1.00 |
| HSPA1A | 0.66 | 0.073 | 1.00 | 0.01 | 0.98 | 1.00 | 0.70 | 0.06 | 1.00 | 0.68 | 0.06 | 1.00 | 0.72 | 0.04 | 0.76 |
| UBB | 0.36 | 0.38 | 1.00 | 0.67 | 0.07 | 1.00 | 0.88 | 3.59E-03 | 6.80E-02 | 0.36 | 0.38 | 1.00 | 0.79 | 0.02 | 0.38 |
| SERPINH1 | 0.46 | 0.25 | 1.00 | 0.66 | 0.08 | 1.00 | -0.22 | 0.60 | 1.00 | 0.53 | 0.18 | 1.00 | 0.27 | 0.51 | 1.00 |
| MRPL18 | 0.47 | 0.24 | 1.00 | 0.62 | 0.10 | 1.00 | -0.03 | 0.94 | 1.00 | 0.50 | 0.21 | 1.00 | 0.55 | 0.15 | 1.00 |
| RPA3 | -0.17 | 0.7 | 1.00 | -0.44 | 0.27 | 1.00 | -0.35 | 0.39 | 1.00 | -0.14 | 0.75 | 1.00 | 0.56 | 0.15 | 1.00 |
| CRYAB | 0.57 | 0.14 | 1.00 | 0.80 | 0.02 | 0.38 | 0.43 | 0.28 | 1.00 | 0.59 | 0.12 | 1.00 | 0.48 | 0.23 | 1.00 |
| CAMK2A | 0.33 | 0.18 | 1.00 | 0.69 | 0.06 | 1.00 | 0.61 | 0.11 | 1.00 | 0.45 | 0.26 | 1.00 | 0.66 | 0.07 | 1.00 |
| CAMK2G | 0.34 | 0.4 | 1.00 | -0.03 | 0.95 | 1.00 | 0.45 | 0.26 | 1.00 | 0.71 | 0.05 | 0.95 | 0.81 | 0.02 | 0.38 |
| MTOR | 0.15 | 0.73 | 1.00 | -0.17 | 0.69 | 1.00 | -0.46 | 0.25 | 1.00 | 0.43 | 0.29 | 1.00 | -0.03 | 0.94 | 1.00 |
| HSPB8 | 0.15 | 0.73 | 1.00 | 0.06 | 0.89 | 1.00 | 0.30 | 0.47 | 1.00 | 0.65 | 0.08 | 1.00 | -0.03 | 0.94 | 1.00 |

**Supplemental Table 5.**
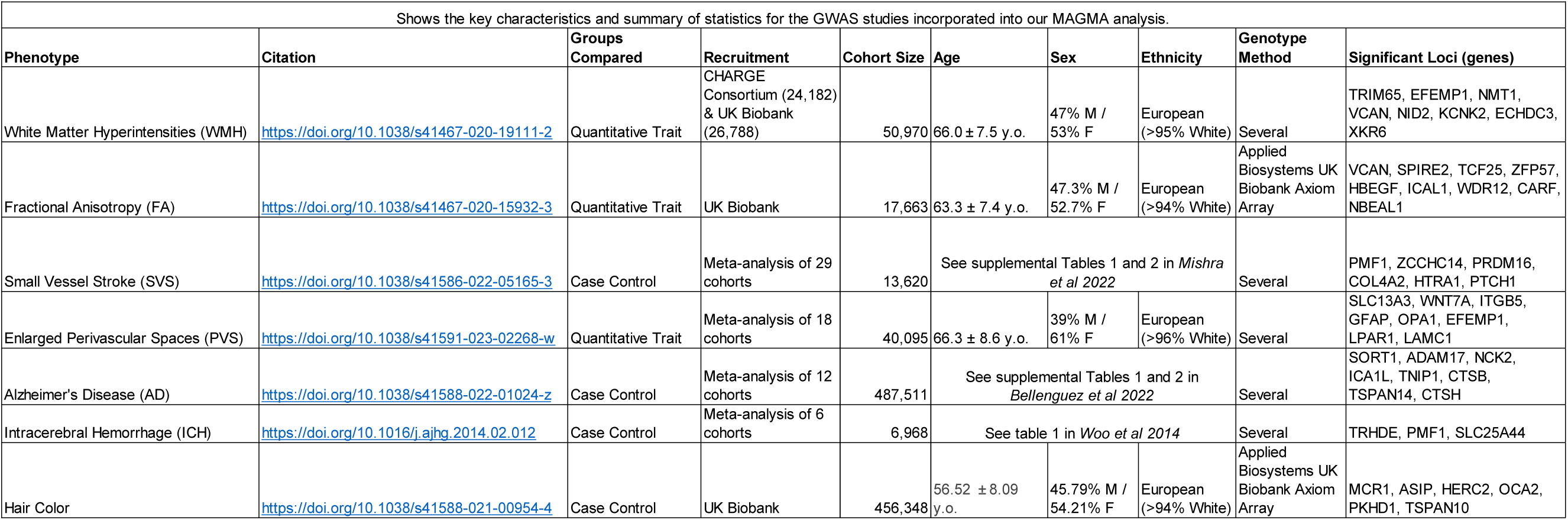
Summary of characteristics for the GWAS used in MAGMA.

